# Dual species transcriptomics reveals core metabolic and immunologic processes in the interaction between primary human neutrophils and *Neisseria gonorrhoeae* strains

**DOI:** 10.1101/2022.02.28.482360

**Authors:** Vonetta L. Edwards, Aimee D. Potter, Adonis D’Mello, Mary C. Gray, Amol C. Shetty, Xuechu Zhao, Katherine M. Hill, Stephanie A. Ragland, Alison K. Criss, Hervé Tettelin

## Abstract

*Neisseria gonorrhoeae* (the gonococcus, Gc) is the causative agent of the sexually transmitted infection gonorrhea. Gc is a prominent threat to human health by causing severe and lifelong clinical sequelae, including infertility and chronic pelvic pain, which is amplified by the emergence of “superbug” strains that are resistant to all current antibiotics. Gc is highly adapted to colonize human mucosal surfaces, where it survives despite initiating a robust inflammatory response and influx of polymorphonuclear leukocytes (PMNs or neutrophils) that typically clear bacteria. Here, dual-species RNA-sequencing (RNA-seq) was used to define Gc and PMN transcriptional profiles alone and after infection. Three strains of Gc and three human donors’ transcriptional responses were assessed to characterize core host and bacterial responses. Comparative analysis of Gc transcripts revealed major overlap between the Gc response to PMNs, iron, and hydrogen peroxide; specifically, the TonB system and TonB dependent transporters (TDT) were upregulated in response to PMNs. We experimentally confirmed that induction of the iron-dependent TDT TbpB is responsive to the presence of PMNs and that *tonB* is required for Gc survival from PMNs. Pathway analysis of PMN transcripts induced by Gc infection revealed differential expression of genes driving pathways involved in cell adhesion and migration, inflammatory responses, and inflammation resolution. Production of pro-inflammatory cytokines, including IL1B and IL8, the adhesion factor ICAM1, and the anti-inflammatory prostaglandin PGE2 was confirmed to be induced in PMNs in response to Gc. Together, this study represents a comprehensive and experimentally validated dual-species transcriptomic analysis of three isolates of Gc and primary human PMNs that gives insight into how this bacterium survives innate immune onslaught to cause disease in humans.

## INTRODUCTION

Gonorrhea, caused by the bacterial pathogen *Neisseria gonorrhoeae* (the gonococcus, Gc), is one of the most common sexually transmitted infections in the United States and globally (1, 2). Drug resistant gonorrhea is an emergent threat to global health, as Gc has acquired resistance to antibiotics including β-lactams, tetracyclines, and fluoroquinolones (3-5). Intramuscular administration of ceftriaxone is the only current frontline antibiotic treatment recommended for uncomplicated gonococcal infection in the United States (6) and high-level ceftriaxone-resistant strains of Gc have been isolated globally (7-10), highlighting the need for new therapeutic approaches for gonorrhea.

Many of the sequelae from gonococcal infection are associated with a sustained inflammatory response featuring polymorphonuclear cells (neutrophils, PMNs). PMNs release proteases, antimicrobial peptides, and reactive oxygen species (ROS), which are delivered to pathogens from cytotoxic granules during phagocytosis and in neutrophil extracellular traps (NETs) (11). However, intact Gc are observed within and attached to PMNs in male urethral exudates and female cervical secretions, and Gc can be cultured from these specimens (12). Gc evades PMN clearance by undergoing antigenic and phase variation to avoid antibody-mediated opsonization, expressing gene products that defend against toxic PMN species, modulating their delivery to phagolysosomes, and delaying the spontaneous apoptosis of PMNs (12-20). However, to date there has not been a comprehensive analysis of the mechanisms used by Gc to avoid and manipulate the PMN response.

It is highly unusual for a pathogen to survive exposure to PMNs. To systematically investigate the mechanisms PMNs direct against Gc, and conversely that Gc uses to resist PMN clearance, we performed dual-species RNA-seq transcriptional profiling of Gc interacting with adherent, IL8 primed primary PMNs from three unrelated individuals. Our study of host and pathogen genes, functions, and pathways associated with Gc-PMN interactions provides the first detailed dual-species transcriptomic analysis of the response of PMNs from multiple donors to multiple Gc strains and vice-versa. This information can be used to inform new therapeutic approaches for drug-resistant infections.

## RESULTS

### Modeling Gc-PMN interactions during human infection

To investigate interactions of Gc with human PMNs, we used an infection model with adherent, IL8 treated primary human PMNs to mimic the tissue-migrated state of PMNs in human disease (21, 22). We used three isolates of Gc: 130 (also known as Opaless), a constitutively piliated, Opa^-^ derivative of strain FA1090 (23); H041 (also known as WHO X), a multidrug-resistant clinical isolate (10); and 3×130, a variant of 130 we engineered that carries 3 antibiotic resistance alleles – *penA, penB* and *mtrR* – from H041 (this study, **see Materials and Methods**). Antibiotic resistance in H041 and an increase in resistance in 3×130 towards multiple antibiotics was verified by measuring Minimum Inhibitory Concentrations (MIC) (**Table S1**). All three isolates of Gc exposed to PMNs displayed an initial decrease in viability at 30 minutes, followed by a recovery and outgrowth period starting at 1 h post infection (**Fig S1**). In comparison, Gc constitutively expressing the CEACAM-binding OpaD exhibited lower initial survival and did not exhibit outgrowth after PMN challenge (**Fig S1**) (23). PMNs remain intact over this time course (24). We examined Gc at the start of the outgrowth period in order to define the transcriptional response of Gc that have resisted PMN killing. The use of 130 and 3×130 avoided potential consequences of Opa phase variation (25).

We investigated the conserved transcriptional responses of 130, 3×130, and H041 and primary PMNs from three unrelated individuals, alone and in co-culture. Within the PMN preparations, neutrophils represented over 95% of the purified cell population (**Fig S2**). Samples were harvested for RNA extraction at the time of Gc addition to PMNs (Gc+PMN_0h), and after 1 h of infection (Gc+PMN_1h). For comparisons, we also included PMNs without infection at 0 h (PMN_0h) and 1 h (PMN_1h), and Gc incubated in media without PMNs for 1 h (Gc_1h) (**Fig 1**). Extraction of high quality and quantity RNA from PMNs is challenging, as neutrophil granules contain RNases that are released upon cell lysis (26). The addition of EDTA during lysis allowed RNA of sufficient quality and quantity to be collected for the preparation of libraries and subsequent sequencing (**SI Appendix**).

**Figure 1:**
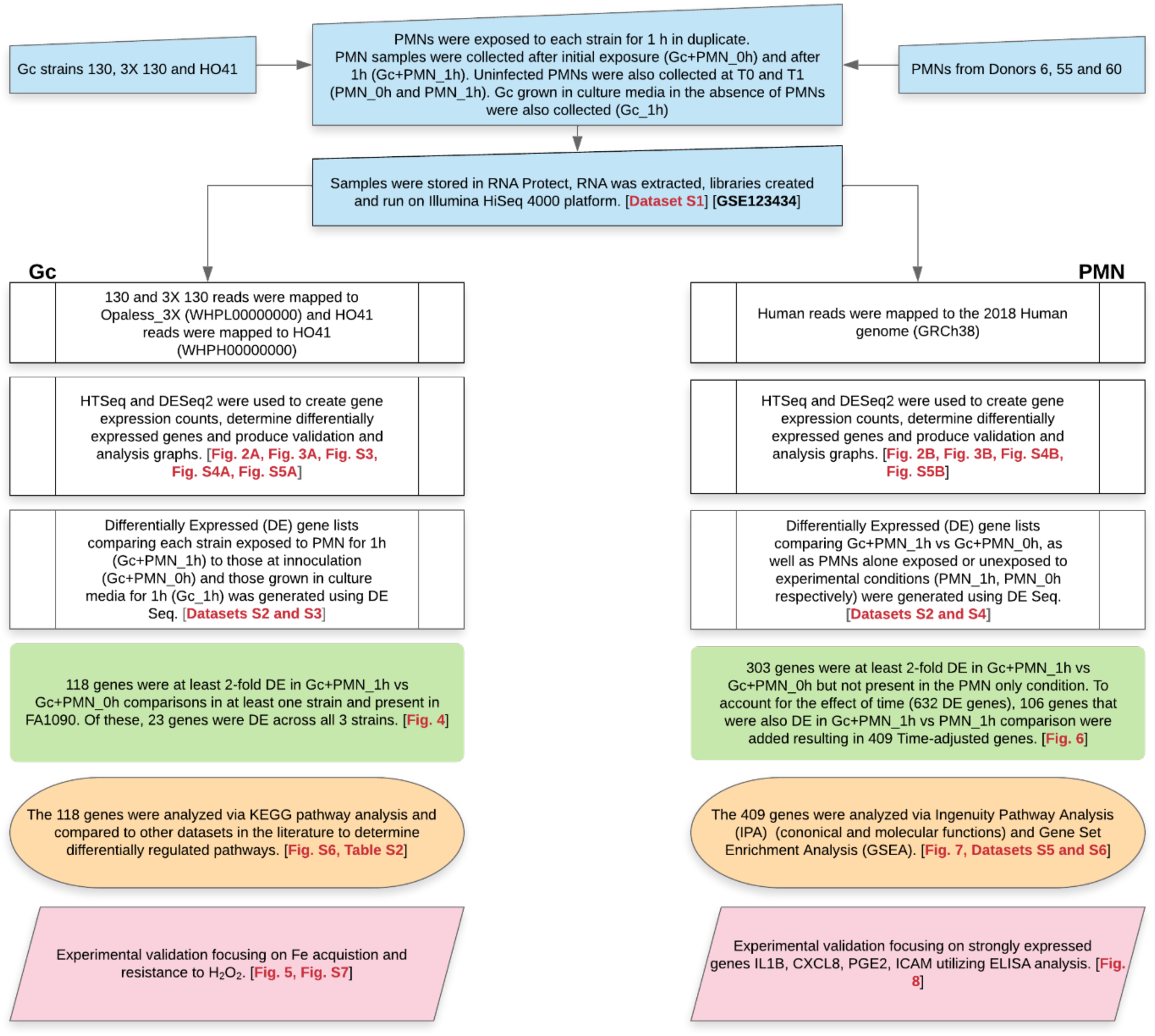
Study overview and associated Figures and Datasets.

### Infection drives consistent differences in gene expression profiles of Gc and human PMN transcriptomes across strain background and PMN donors

In order to compare gene expression across Gc strains, a PanOCT analysis of orthologous genes (27) was performed between the FA1090 (used for annotation purposes), 3×130, and H041 genome sequences (**Fig S3**). From the 1621 PanOCT clusters of orthologs identified, clusters harboring multiple genes from the same genome (e.g. paralogs or gene fragments) were filtered out, resulting in 1576 clusters of core genes used for combined transcriptome analysis (**Dataset S1**).

A major challenge in simultaneous dual RNA-seq profiling of a host and pathogen is obtaining sufficient sequence coverage for both organisms, where mammalian cell transcripts can be 100 times more abundant than bacterial transcripts (28). To ensure sufficient numbers of sequencing reads were obtained, we generated saturation curves of reads mapped to the Gc and human genomes (**Fig 2**). All samples plateaued or approached plateau, indicating that sequencing depth was adequate. The minimum required mapped reads to reach saturation for the ∼1600 core protein coding genes in Gc was 150,000-200,000. For host transcripts, the minimum required mapped reads to reach saturation for the ∼17,500 genes in the human genome was 2,500,000. The proportion of Gc to total mapped transcripts in infected PMN samples was ∼25-35%. A complete overview of sequenced and mapped reads is provided in **Dataset S1**.

**Figure 2:**
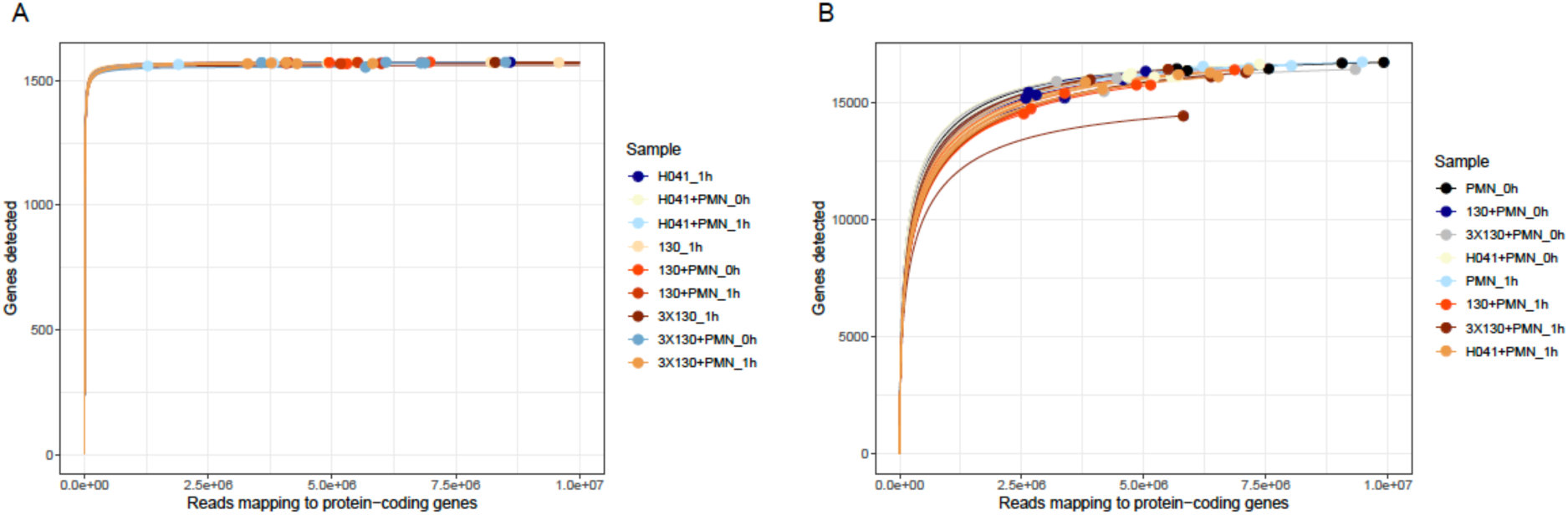
Rarefaction curves of RNA-seq reads mapped to Gc and Human genes. A) Rarefaction curves of reads mapped to the Gc core genes (1621 genes, Dataset S3) for samples containing 130, 3×130, and H041 reads. B) Rarefaction curves of reads for all PMN containing samples mapped to human genes. Samples with curves that plateau have sufficient reads mapped across the respective genome to achieve saturation.

We observed distinct expression profiles for Gc core genes by principal component analysis (PCA) (**Dataset S2, Fig 3A**). PC1 (36.51% of variation) separated samples by experimental condition, indicating that the majority of the Gc response to PMNs is strain-independent. PC2 (22.03%) separated samples by strain background, with 130 and 3×130, which are genetically more closely related, grouping more closely than with H041. A bootstrapped dendrogram of Gc-containing samples also showed grouping by strain background (**Fig S4**). Similarly, PCA was performed on PMN samples (**Fig 3B**). PC1 accounted for 31% of the variation and was associated with time of incubation (1h) vs (0h). This result indicates that the transcriptional response of PMNs is primarily impacted by culture *ex vivo*. Donor-specific responses in PC2 accounted for only 8.8% of the variation between samples, suggesting that PMNs behaved similarly across experimental conditions. This was also reflected in a bootstrapped dendrogram of PMN-containing samples (**Fig S4**). The remaining PCs accounted for less than 10% of the variation between samples for both Gc and PMNs (**Fig S5A**).

**Figure 3:**
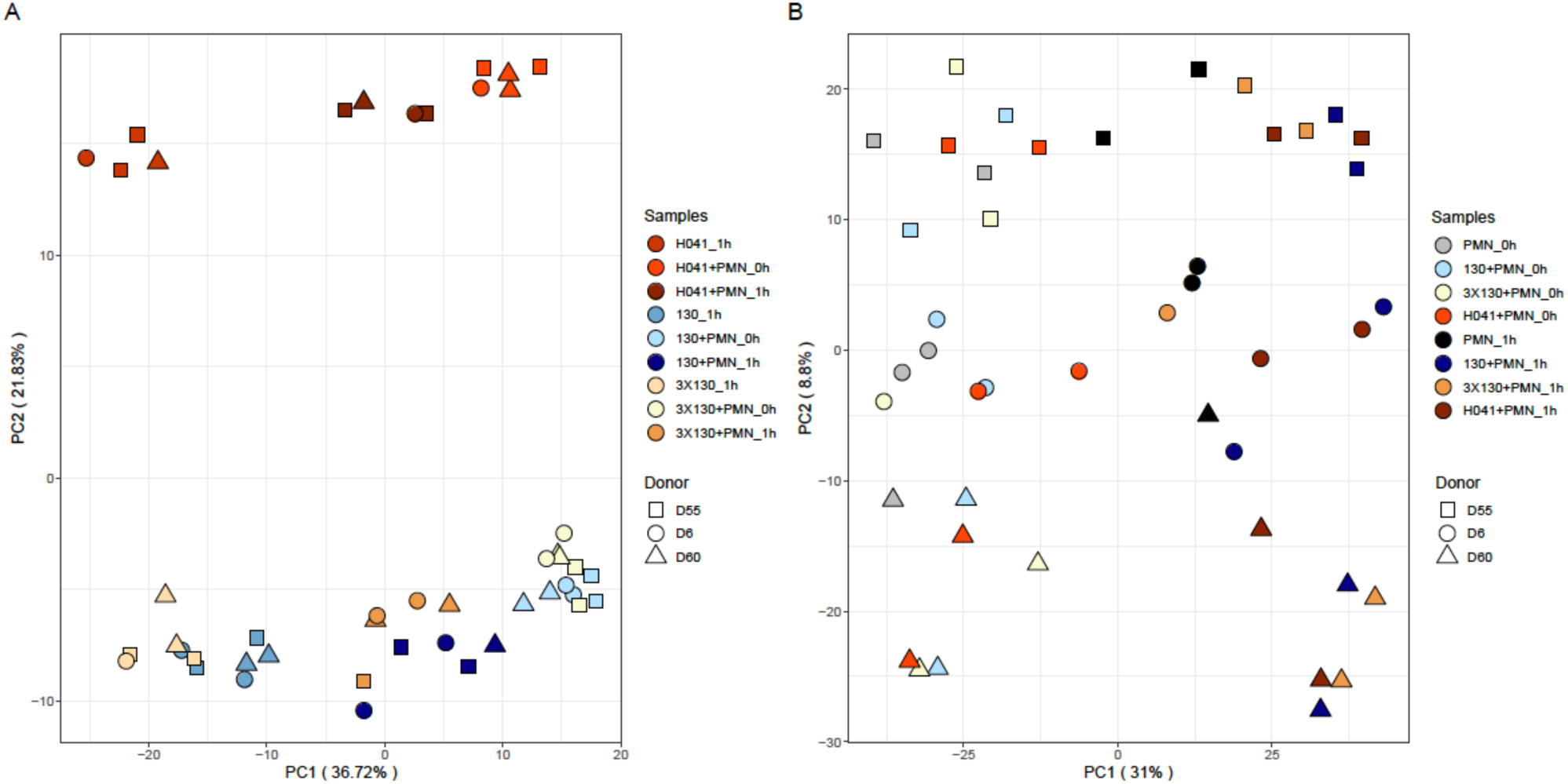
Principal component analysis of Gc and PMNs alone and after infection. PCA of core gene expression profiles for Gc (A) and PMNs (B). Shapes represent PMN donors. Colors represent Gc strains. Light and dark shades represent the 0h and 1h infection conditions, respectively.

### Gc differentially regulates selected gene subsets in response to infection

Comparisons of PMN-exposed Gc (Gc+PMN_1h) to time of inoculation (Gc+PMN_0h) for each strain revealed 80 differentially expressed (DE) genes for 130, 165 for 3×130, and 60 for H041 are visualized in an Upset plot (Green bars **Fig 4A and B, Dataset S3**). The individual or connected dots in the Upset plot represent the various intersections of genes that were either unique to, or shared among comparisons, comparable to a Venn diagram. Vertical bars constitute the number of DE genes unique to specific intersections. Horizontal bars constitute the number of DE genes for each specific comparison. Genes from these lists were cross-referenced with clusters of core genes to define 118 core genes as DE (FDR ≤ 0.05 and log_2_(fold change) ≥ 1 or ≤ -1) in at least one strain (**Fig 1, Fig 4C, Dataset S3**). Of the 118 Gc DE genes, 23 genes were significantly up- or down-regulated in the same direction among all three Gc isolates (Gc+PMN_1h vs Gc+PMN_0h) (**Blue bar Fig 4C, and Fig 4D**). We also compared PMN-exposed Gc vs. unexposed Gc at 1 h (Gc+PMN_1h vs Gc_1h) for each of the 118 genes to contextualize this list and account for the pronounced physiological effect of incubation in culture media (**Dataset S3, Fig 3A, Fig 4**). The 118 genes DE by Gc in response to adherent PMNs were compared to RNA-seq datasets representing a variety of environmental factors including iron (29), hydrogen peroxide (H_2_O_2_) (30), anaerobic growth (31), exposure to PMNs in suspension (32), and natural human infection (33) (**Dataset S3**). The Gc response to PMNs overlaps substantially with the anaerobic, H_2_O_2_, and iron responses with 27, 41, and 27 of the 118 DE genes identified, respectively (**Dataset S3**).

**Figure 4:**
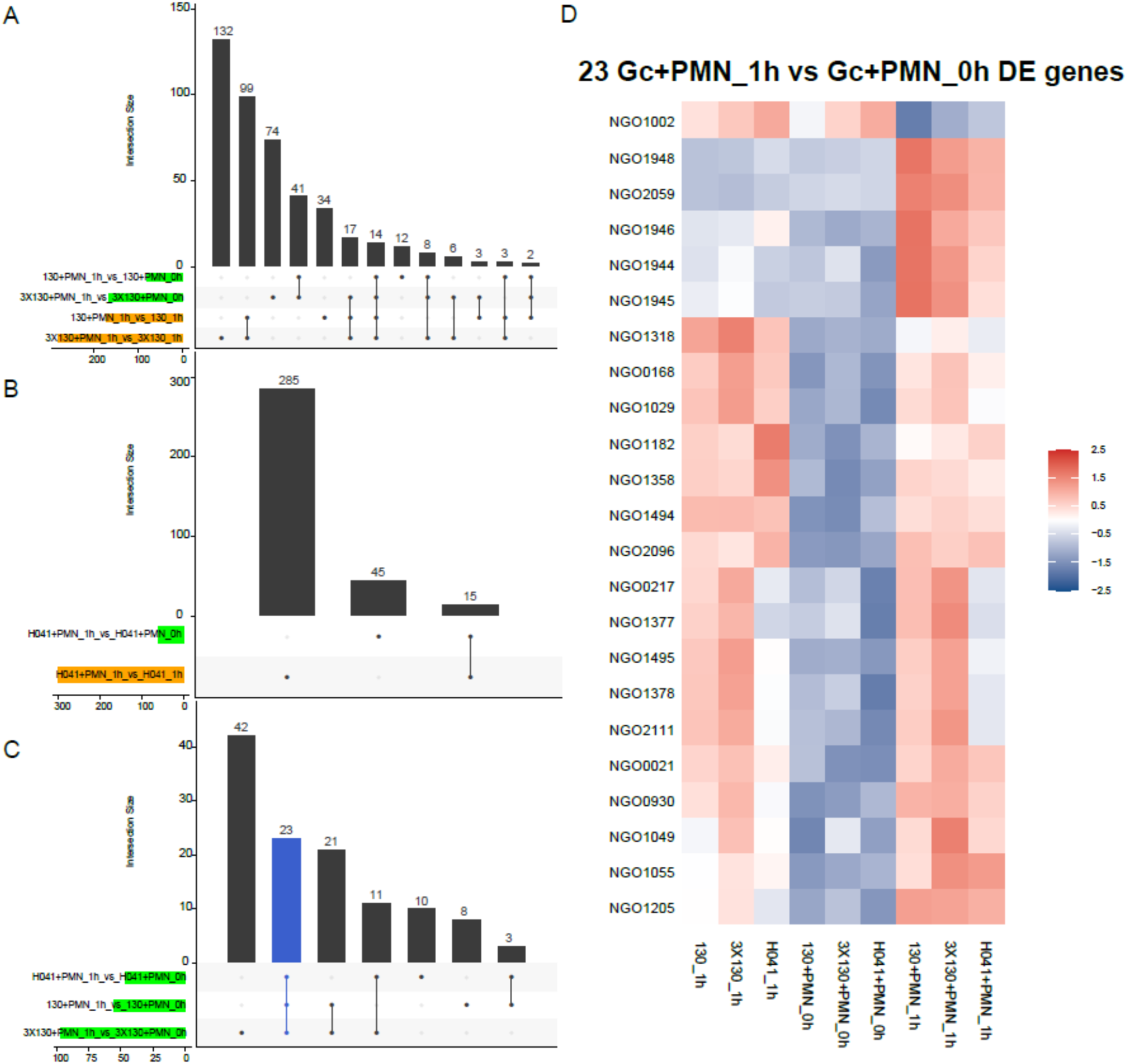
Gc DE gene analysis. A) Upset plot of all (3X)130+PMN_1h vs (3X)130+PMN_0h and (3X)130+PMN_1h vs (3X)130_1h comparisons using all (3X)130 genes. B) Upset plot of H041+PMN_1h vs H041+PMN_0h and H041+PMN_1h vs H041_1h comparisons using all H041 genes. C) Upset plot of all Gc+PMN_1h vs Gc+PMN_0h comparisons using only core genes without paralogs or fragmented genes (from the master list of PanOCT orthologs listed in **Dataset S3 Tab7**). The blue vertical bar is the shared intersection of DE genes for all three strains. D) Z-score heatmap of 23 DE genes (blue bar from panel C).

#### Pathway-centric analysis of Gc DE genes upon exposure to PMNs

Kyoto Encyclopedia of Genes and Genomes (KEGG) pathway analysis (34) of the 118 genes DE in Gc upon exposure to PMNs identified several enriched pathways. Specifically, 83% of Gc genes (5 of 6) annotated for nitrogen metabolism (KEGG pathway 00910) were DE (**Table S2, Fig S6**). These genes can be separated broadly into two categories, nitrogen utilization: carbonic anhydrase (*cah*, NGO0574), glutamate dehydrogenase (NGO1358), and glutamine synthetase (NGO1600); and anaerobic respiration on nitrite: nitrite reductase (*aniA*, NGO1276) and nitronate monooxygenase (NGO1024). Although not included in KEGG nitrogen metabolism pathway, nitrogen utilization genes glutamyl tRNA amidotransferase (*gatB*, NGO2133) and acetylglutamate kinase (NGO0844), as well as the anaerobically induced nitrogen regulatory protein II (NGO1182), were also DE in Gc exposed to PMNs (**Dataset S3**). While these genes were significantly upregulated in PMN-exposed Gc at 1 h compared to 0h (Gc+PMN_1h vs Gc+PMN_0h), they were also generally downregulated in PMN-exposed Gc compared to unexposed Gc (Gc+PMN_1h vs Gc_1h), suggesting that nitrogen metabolism is stimulated during Gc culture with PMNs (**Dataset S3**). This expression pattern was most prominent for *aniA* (NGO1276) (**Dataset S3, Fig 4D**). The nitrite reductase, *aniA*, is induced in response to anaerobic culture and is required for Gc anaerobic growth (31, 35). Stimulation of nitrogen metabolism in Gc may facilitate production of nitrite, which is used by Gc as an electron acceptor under anaerobic conditions. This expression pattern suggests that Gc may also experience oxygen-depleted conditions when exposed to PMNs, possibly due to PMN oxygen consumption, and explain, in part, the overlap of our dataset with the anaerobic stimulon (**Dataset S3**) (31).

KEGG analysis of Gc genes that are DE upon exposure to PMNs also revealed enrichment of ABC transporters (KEGG pathway 02010): *fbpA* (NGO0217), *potF4* (NGO1494), *cysAW* (NGO0445-NGO0446), and *tcyABC* (NGO0372-NGO0374) (**Table S2**). Manual inspection of DE genes revealed additional ABC transporters, currently unannotated in KEGG (NGO2011-NGO2014). In total, ABC transporters comprise the largest subset of the 31 genes downregulated in Gc in response to PMNs (10 of 31). These included three related complexes: a cystine transporter (NGO2011-NGO2014), cysteine transporter (*tcyABC*, NGO0372-NGO0374), and sulfite/thiosulfate transporter (*cysAW*, NGO0445-NGO0446), as well as cysteine synthase A (NGO0340) (36, 37). Cysteine, either imported or synthesized from sulfite or cystine, is required for the synthesis of glutathione, which confers resistance to oxidative stress. Gc produces high concentrations of glutathione compared to other bacteria (>15mM) and does not encode glutathione importers (38). As such, Gc production of cysteine is thought to be important for the high level oxidative stress resistance of Gc (37). While genes involved in cysteine transport and production were significantly downregulated in PMN-exposed Gc at 1h compared to 0h (Gc+PMN_1h vs Gc+PMN_0h), they were upregulated in PMN-exposed Gc compared to unexposed Gc (Gc+PMN_1h vs Gc_1h) (**Dataset S3**). These results imply that import of sulfite, cystine, and cysteine is an important component of Gc response to PMNs.

#### Gc upregulates oxidative stress response genes upon exposure to PMNs

Of the 118 Gc genes that were DE upon exposure to PMNs (Gc+PMN_1h vs Gc+PMN_0hr), 41 were also reported to be responsive to H_2_O_2_ (**Dataset S3**) (30). NGO2059 (*msrAB*), encoding Methionine Sulfoxide Reductase polyprotein (MsrA/B), exhibited the greatest fold change upon exposure to PMNs (**Dataset S3**). MsrA/B is surface-exposed and conserved among Gc strains. It is associated with repair of oxidative damage in Gc and is required for survival from oxidative stressors such as H_2_O_2_ and superoxide anions (39). *MsrAB* is known to be regulated in part by the alternative sigma factor Ecf (NGO1944), which also auto-upregulates a predicted five gene operon (NGO1944, NGO1945, NGO1946, NGO1947, and NGO1948) (40). *Ecf, msrAB*, and all members of the *ecf* operon except NGO1947, were upregulated in Gc exposed to PMNs in all three Gc strains (**Fig 4D**) (40).

In addition to *msrAB*, transcripts for other H_2_O_2_ responsive genes are also more abundant in Gc exposed to PMNs (**Dataset S3**). For example, NGO1767, encoding catalase (*kat*) which catalyzes the decomposition of H_2_O_2_ to oxygen and water and is required for Gc survival from H_2_O_2_, was also upregulated in Gc exposed to PMNs (41). Eighteen of the 41 identified H_2_O_2_ responsive genes are also responsive to iron and are discussed in depth below (NGO0025-*mpeR*, NGO0217-*fbpA*, NGO0554, NGO0655, NGO0863, NGO0929-*metF*, NGO1029-*fumC*, NGO1276-*aniA*, NGO1318-*ccmA*, NGO1377-1379-*tonB*/*exbB*/*exbD*, NGO1495-1496-*tbpAB*, NGO1769-*ccpR*, NGO1779, NGO2092, and NGO2093-*fetA*). An additional 12 are implicated in nutrient transport (NGO1049, NGO1205-*tdfJ*, NGO1494-*potF4*, NGO2050-*efeO*, NGO2109-*hpuB*) or metabolism (NGO0848, NGO0918, NGO0919, NGO0956, NGO1294, NGO1931, NGO2133). Nutrient homeostasis, particularly nutrient metals, can affect oxidative stress resistance (14). The remaining 7 non-iron responsive genes identified are largely genes of unknown function (NGO0489, NGO0568, NGO0775, NGO1002-*traA*, NGO1055-*yciA*, NGO1566, NGO1588). Gene products that have been shown to defend against oxidative stress that are responsive to PMN exposure, but did not overlap with the H_2_O_2_ responsive genes identified by Quillin et. al (30), included Cytochrome C peroxidase (*ccpR*, NGO1769) (14) and lactate permease (*lctP*, NGO1449) (42).

Based on these results, we hypothesized that Gc exposure to PMNs primes an oxidative stress response that enhances bacterial survival from ROS. To test this, Gc strain 130 was exposed to PMNs for 1h or exposed to media alone, then treated with increasing concentrations of H_2_O_2_ for 15 minutes. Exposure to PMNs modestly (∼12.6%) but significantly increased the percent of Gc recovered after H_2_O_2_ treatment across a range of concentrations, indicating that the transcriptional program initiated by PMN challenge helps defend against oxidative stress (**Fig S7**).

#### Differential expression of metal acquisition systems indicates ways Gc overcomes nutritional immunity when exposed to PMNs

Ten of the 23 DE genes shared by all 3 Gc strains were metal transporters, and 27 of the 118 total DE genes identified in any Gc strain were associated with iron acquisition (**Fig 4D, Dataset S3**). These results led us to hypothesize that Gc responds to PMN challenge by increasing expression of iron acquisition proteins to overcome human nutritional immunity. We focused on the TonB-dependent transferrin receptor TbpAB (NGO1495-NGO1496), which is required for symptomatic infection in male urethral challenge and shows Fur-dependent repression of expression in high iron conditions (29, 43). Expression of *tbpB* was induced following incubation with PMNs (Gc+PMN_1h vs Gc+ PMN_0h). In line with the RNA-seq data, TbpB protein levels in strain 130 significantly increased after 1 h of PMN infection (Gc+PMN_1h vs Gc+PMN_0h). Strikingly, at both the RNA and protein level, TbpB expression was lower in Gc exposed to PMNs after 1h (Gc+PMN_1h) compared to those that were not (Gc_1h) (**Fig 5A, B**). This observation led us to investigate if this phenomenon was observed for other Gc metal acquisition genes. In addition to TbpAB, these include genes encoding the TonB/ExbB/ExbD system (NGO1377-1379), which powers import of metals via outer membrane transporters; FetA (NGO2093), which acquires iron from siderophores produced by other bacteria; FbpA (NGO0217), a periplasmic metal binding protein for ferric iron; HemO (NGO1318), a heme oxygenase required for iron utilization from heme (44); NGO2111, a homolog of *N. meningitidis* Slam2 (NMB1917) required for translocating the hemoglobin transporter component HpuA to the bacterial surface (45); and NGO0554, a predicted hemophore (46) that is required for high level resistance to H_2_O_2_ and is repressed by iron (19). Although not overlapping with the iron-responsive regulon identified by Yu et. al (29), other genes DE in Gc after exposure to PMNs included ZnuA (NGO0168) and ZnuC (NGO0170), which with ZnuB form a transport system for zinc and manganese from the periplasm into the cytoplasm (47, 48); NGO1049, a predicted periplasmic metal transport protein (NGO1049) (49); EfeO (NGO2050), a putative periplasmic ferrous iron transporter; and the TonB-dependent, iron-regulated transporters HpuB (NGO2109), which imports hemoglobin, TdfJ (NGO1205), which acquires zinc from psoriasin (50, 51), and TdfF (NGO0021), whose ligand is unknown but is important for survival of Gc inside epithelial cells (52). Similar to the expression pattern observed for TbpB, the iron-responsive genes were most dramatically DE in the Gc_1h condition relative to Gc+PMN_0h, with a less potent response in the Gc+PMN_1h condition (**Dataset S3**). This overall expression pattern was likely a driver of the distribution of conditions along PC1 in **Fig 2**. The induction of metal acquisition genes in the Gc_1h condition may be explained by the lack of accessible iron for Gc in the infection medium (RPMI+10% FBS), as Gc TbpAB can only use human transferrin, not bovine (53, 54). The dampening of the Gc metal responsive gene expression during exposure to PMNs suggest that nutrient metals, specifically iron, are more available and/or Gc requirements for metals are lower following Gc co-culture with PMNs compared with unexposed Gc. To test the latter possibility, we assessed survival of a Gc *tonB* mutant upon exposure to PMNs, which is incapable of growth on transferrin as a sole iron source (55). Compared to 130, an isogenic Δ*tonB* mutant was significantly reduced for survival at 30, 60, and 120 minutes post-PMN infection (**Fig 5C**), suggesting that TonB-dependent metal acquisition contributes to Gc survival during PMN challenge. In total, these results show a coordinated transcriptional response of Gc towards PMNs, particularly in oxidative stress responses and nutrient acquisition.

**Figure 5:**
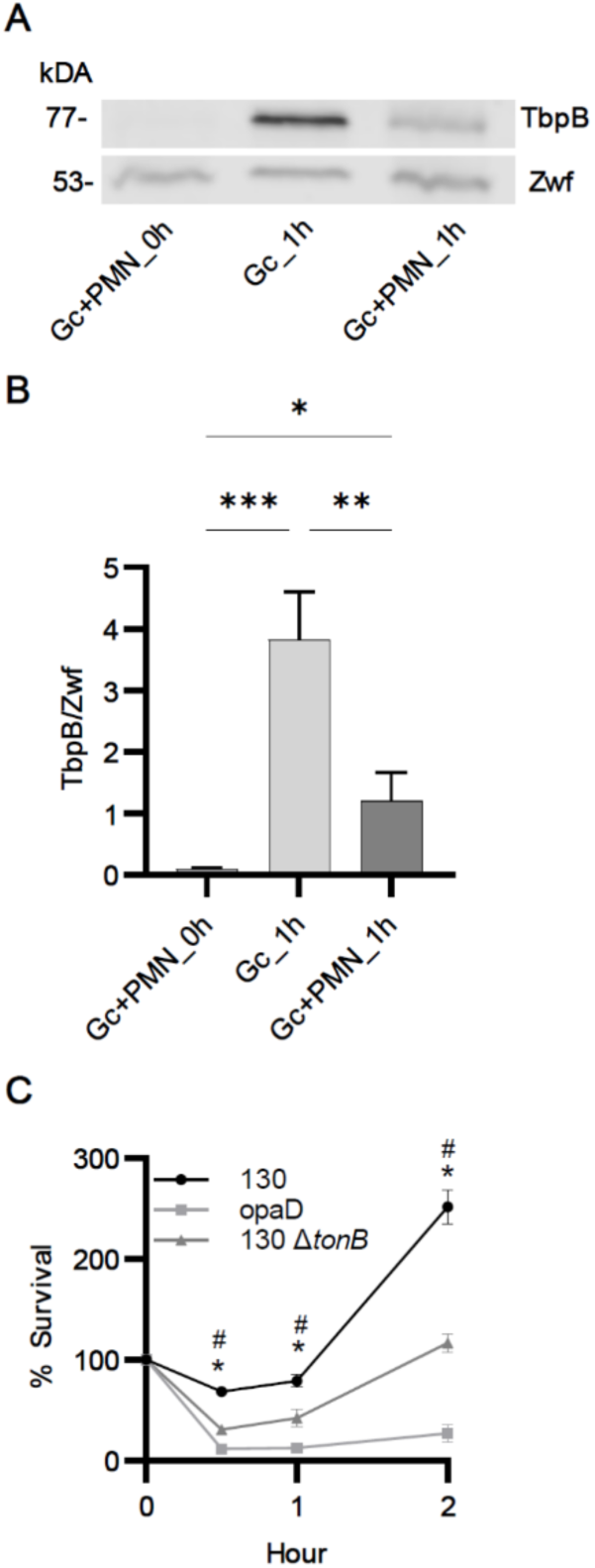
Increased production of TonB-dependent metal acquisition systems contribute to Gc survival from PMNs. A) Gc strain 130 was inoculated onto IL8-treated, adherent human PMNs or into media without PMNs, and incubated for 1h. PMNs were treated with saponin, and Gc were collected and processed for Western blotting. Gc lysates were separated by 10% SDS-PAGE, transferred to a nitrocellulose membrane, and stained with rabbit anti-TbpB polyclonal antisera. The intensity of TbpB is reported relative to the loading control Zwf, which was recognized with rabbit anti-Zwf antiserum. A representative blot from a single experiment is shown. B) Quantification of TbpB/Zwf ratio from 3 independent experiments, using PMNs from 3 unrelated individuals. Error bars indicate SD. Significance was determined by one-way ANOVA with Holm– Sidak correction for multiple comparisons, * indicates *p* < 0.05, ** *p* < 0.01 and *** *p* < 0.001. C) Strain 130, its isogenic Δ*tonB* mutant, or another isogenic mutant constitutively expressing OpaD were exposed to adherent, IL8-treated primary human PMNs. Percent Gc survival was calculated by enumerating colony-forming unit (CFU) from PMN lysates at 30, 60, and 120 min and reported as the percent of CFU for that strain at 0 min. Significance was determined by two-way ANOVA with Holm–Sidak correction for multiple comparisons, * and ^#^ indicate *p* < 0.05 for 130 Δ*tonB* vs 130 and OpaD vs 130, respectively. *n* = 3 independent experiments.

### PMNs initiate an inflammatory and migratory response to Gc infection

To complement the analysis of Gc responses to PMNs, we examined the PMN responses to Gc that were shared across the three PMN donors. PMN genes that were DE after exposure to Gc for 1 h compared to 0 h (Gc+PMN_1h vs Gc+PMN_0h) were identified (FDR ≤ 0.05 and log_2_(fold change) ≥ 1 or ≤ -1). There were 1916 DE genes in 130-exposed PMNs, 1420 in 3×130-exposed PMNs, and 1644 in H041-exposed PMNs (**Fig 6A, Dataset S4**). In comparison, 1677 genes were DE in PMNs incubated 1 h in media without Gc (PMN_1h vs PMN_0h); these reflect infection-independent PMN DE genes (**Fig 6A, Dataset S4**). We then determined the shared PMN response to Gc, defined as the 303 PMN genes that were at least 2-fold DE in the same direction upon exposure to all 3 Gc backgrounds (**Fig 6A - green bar**). To capture genes with magnitude differences during infection but also changing with time, 632 time-dependent genes (common to all four 1 h vs 0 h comparisons, **Fig 6A - purple bar**) were intersected with DE genes in Gc+PMN_1h vs PMN_1h comparisons (**Fig 6B**). The resulting 106 genes (**Fig 6B - green bar**) were added to the 303 genes that were DE for PMNs infected with Gc over 1 h (Gc+PMN_1h vs Gc+PMN_0h), yielding the Time-Adjusted PMN list of 409 DE genes (**Fig 1A and 1B – sum of green bars, Fig 6, Dataset S4, Fig S5**). Approximately 50% of the Time-Adjusted PMN genes had a four-fold or more increase across all 3 strains. 397 genes were upregulated, while 12 genes were downregulated.

**Figure 6:**
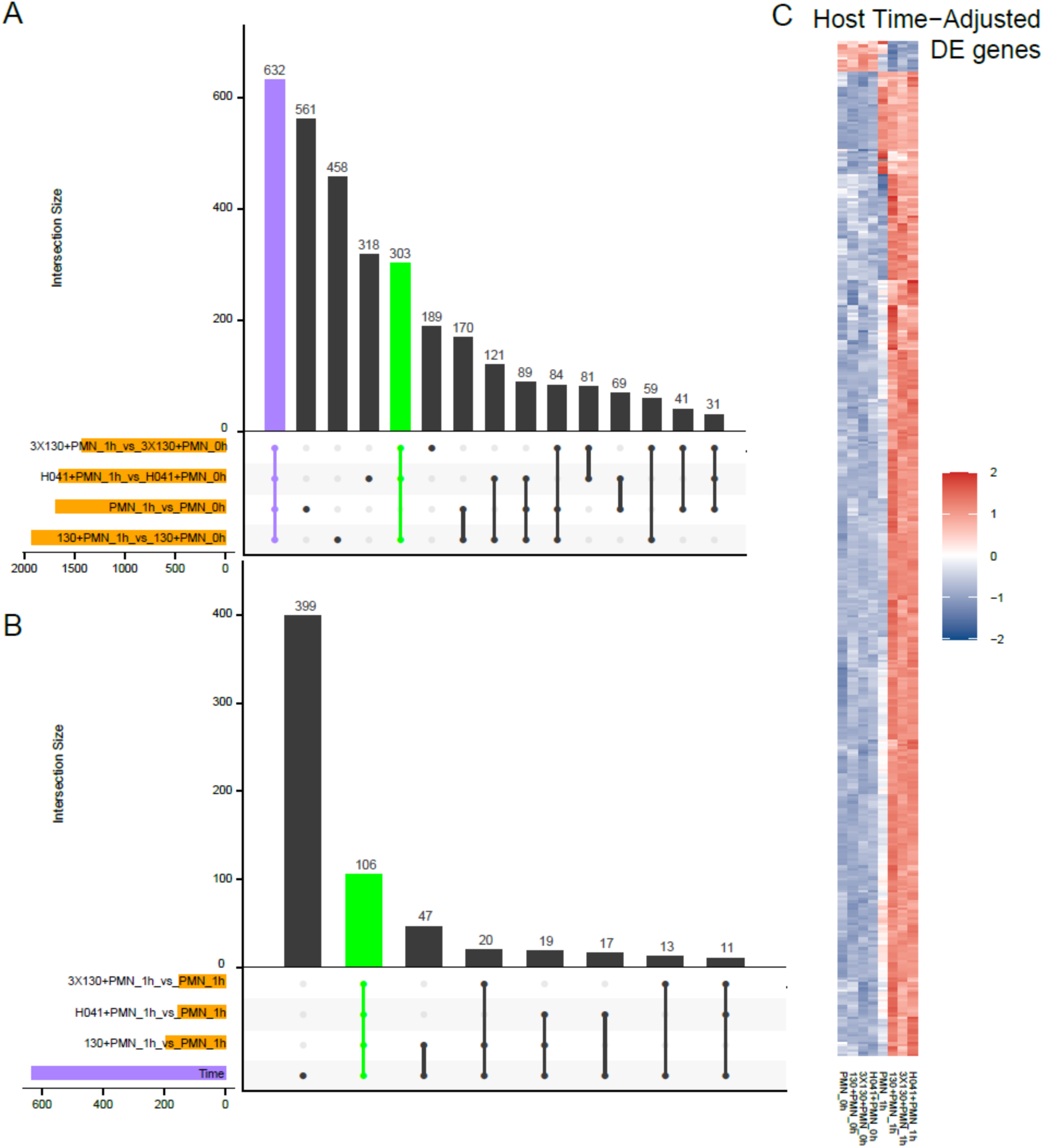
Identification of PMN genes that are DE over time with Gc. **A**) Upset plot of PMN DE genes for all PMN_1h vs PMN_0h comparisons, with and without Gc infection. The green vertical bar indicates 303 genes that are DE in all three +Gc conditions. Purple bars indicate 632 genes that are DE in all conditions of 1h vs 0h regardless of Gc infection, i.e. time-dependent. B) To capture genes with magnitude differences during infection and also changing with time, 632 time-dependent genes (purple horizontal bar transposed from panel A) were intersected with DE genes in Gc+PMN_1h vs PMN_1h comparisons, yielding 106 DE genes. These 106 genes (green vertical bar) were added to the 303 DE genes in A to generate the Time-Adjusted list of 409 DE genes. C) Z-score heatmap of the 409 Time-Adjusted DE genes (red is upregulated, blue is downregulated).

#### Pathway-centric analysis of PMN DE genes upon exposure to Gc – Canonical Pathways and Molecular Functions

The 409 Time-Adjusted PMN DE genes were subjected to Ingenuity Pathway Analysis (IPA). Since pathways specific to PMNs are underrepresented in IPA, all primary cell subsets in the IPA database were used for analysis to identify significantly enriched Canonical Pathways and Molecular Functions (**Dataset S5**). We selected Molecular Functions for further analysis as these better represented activities associated with neutrophils, were divided into specific subcategories, and included pathways important for inflammation, cellular signaling, and cellular defense from infection. As such, we focused on the top 20 (5%) most highly expressed time-adjusted PMN DE genes that drove the top 15 most enriched Molecular Functions (**Fig 7, Dataset S4, Dataset S5**).

**Figure 7:**
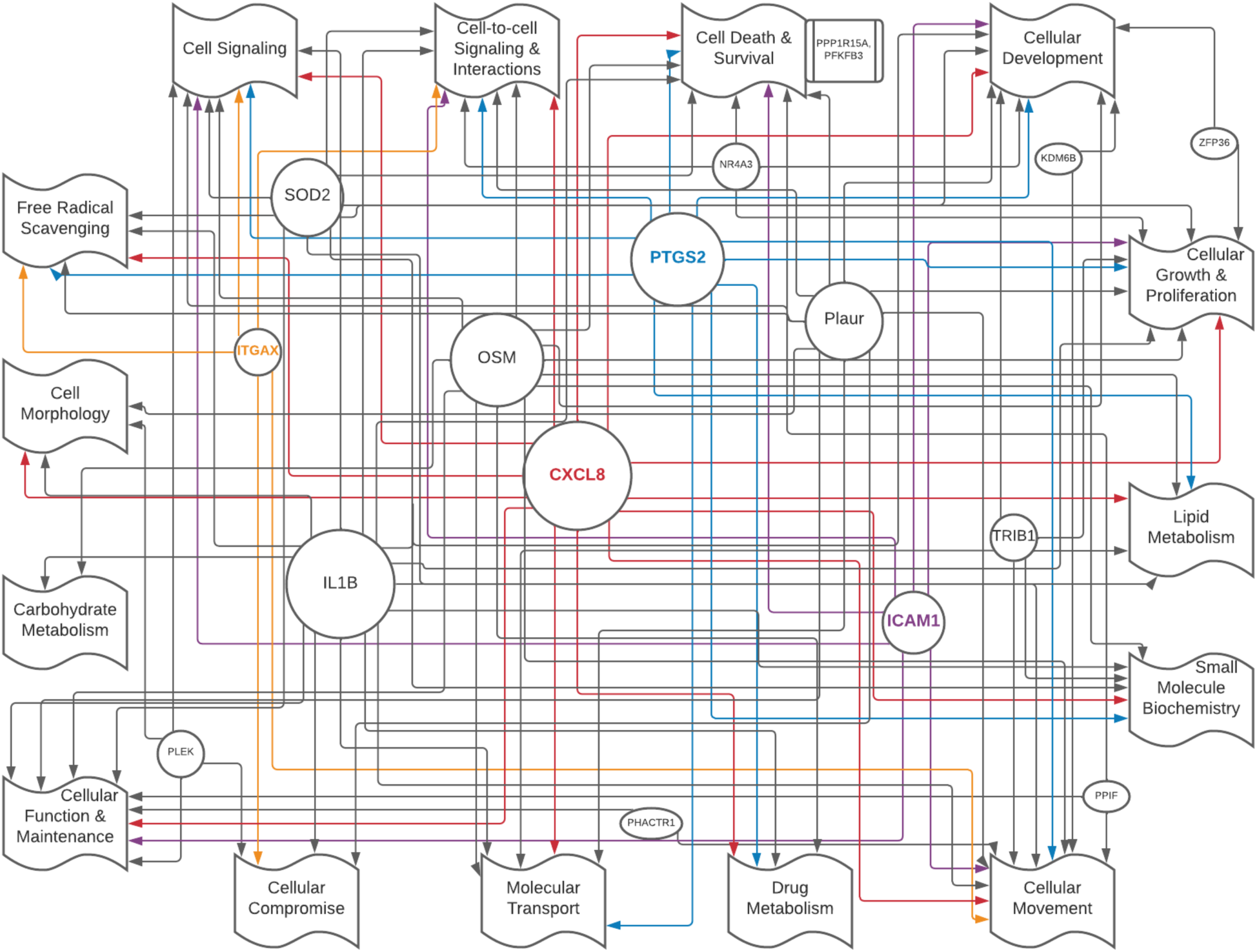
Molecular Functions enriched in time-adjusted PMN genes. Intersection of top 15 IPA Molecular Functions that are significantly enriched (z-score >|2|) and top 20 most highly expressed genes from the 409 time-adjusted PMN genes (see Fig 6C). Genes are linked by arrow(s) to the assigned Molecular Function(s), and the size of the shape reflects the number of associated Molecular Functions. The proteins encoded by the four colored genes or their products are examined in Figure 8. Genes associated with a single Molecular Function are listed in the boxes adjacent to the Molecular Function.

#### PMNs upregulate cytokine secretion and migratory responses

Cell Signaling - Cytokine and Chemokine Related Signaling Pathways was the most enriched subset of Molecular Functions identified by IPA (-log(p-value 13.04)) (**Fig 7, Dataset S5**). Enrichment of these pathways is consistent with one of the hallmarks of Gc infection: robust recruitment of PMNs by pro-inflammatory cytokines to the infection site (12). Analysis of the genes driving enrichment of these two pathways revealed that they encompass a multitude of cytokine transcripts that are upregulated in PMNs following co-culture with Gc (**Fig 7, Dataset S5**). These genes include classical chemokines that PMNs are known to produce upon activation for the purpose of recruiting additional PMNs as well as other leukocytes, including CCL3 (MIP1A), CCL20 (MIP3A), CCL4 (MIP1B), CXCL1 (GRO), CXCL2 (MIP2A), CXCL3 (MIP2B), CXCL8 (IL8), LIF, and VEGFA (56, 57), in addition to less well characterized chemokines CCL3L3 and CCL4L1 (56, 58). We also observed upregulation of genes for pro-inflammatory cytokines IL1A and IL1B, as well as the receptors IL1-R2 and IL1-RN, with IL1B being the most frequent gene driving Molecular Function enrichment (**Fig 7**). IL1 signaling contributes to PMN recruitment, granulopoiesis, degranulation, and NET formation through both direct actions on PMNs and other cell types that subsequently recruit and activate PMNs (59). To characterize the inflammatory environment during Gc-PMN coculture, we conducted multiplexed cytokine profiling of Gc and PMNs at 4 h post-infection, to account for de novo transcription and translation and allow for accumulation of cytokines in the supernatant (**Fig 8A**). Upregulated cytokine transcripts correlated with increased presence of encoded cytokines. Relative to the uninfected control, Gc-infected PMNs released over 4 times (L2FC > 2) the levels of classical cytokines including IL1A, IL1B, IL1RA, IL6, IL8, MIP1A (CCL3), MIP1B (CCL4), CXCL1 (GRO), TNFA, and VEGFA (**Fig 8A**). These results highlight the induction of a robust migratory response of PMNs towards Gc during infection.

**Figure 8:**
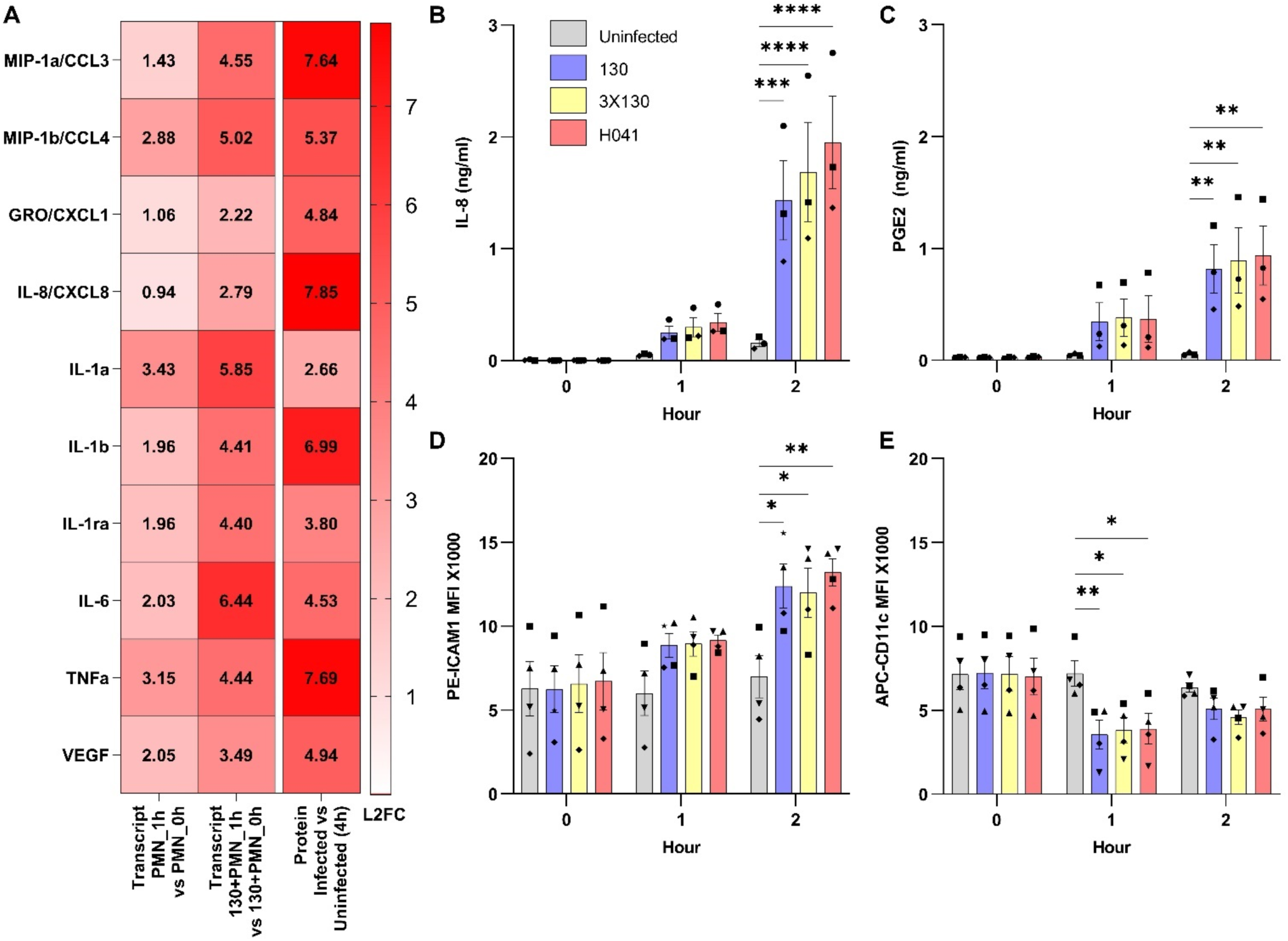
PMNs increase production of pro-inflammatory cytokines and adhesion molecules upon Gc infection. A) Supernatants were collected from uninfected and 130-infected adherent PMNs after 4h, and quantified by multiplex cytokine analysis. Selected targets identified from the 409 Time-Adjusted PMN DE genes are displayed. The Log_2_(Fold Change) (L2FC) in detected protein for 130 infected PMNs at 4h vs uninfected PMNs at 4h is reported. The corresponding L2FC transcript levels identified by RNA-seq for PMN_1h vs PMN_0h and 130+PMN_1h vs 130+PMN_0h is also indicated. B-C) Supernatants were collected from PMNs left uninfected, or exposed to 130, 3×130, or H041 at the indicated timepoints. IL8 (B) and PGE2 (C) concentrations were measured by ELISA. Bars represent the mean ± SEM. *n* = 3 independent experiments. D and E) PMNs were collected and assessed by flow cytometry at the indicated timepoints. ICAM1 (D) and ITGAX (CD11c) (E) surface expression was assessed by Median Fluorescent Intensity (MFI) of PE-ICAM1 and APC-CD11c on CD11b^+^/CD14^-^/CD16^+^/CD49^-^ cells (i.e., neutrophils). Bars represent the mean MFI ± SEM. *n* = 4 independent experiments. B-E) Shapes are data points from different donors’ PMNs. Significance was determined by two-way ANOVA with Holm–Sidak correction for multiple comparisons., *indicates *p* < 0.05.

The chemotactic cytokine CXCL8 (IL8) was one of the most frequent genes driving Molecular Function enrichment, being a part of 13 of the 15 top functions (**Fig 7, Dataset S4**). In PMNs, IL8 can induce upregulation of adhesion factors, priming of the oxidative burst, and release of lysosomal enzymes, and PMNs have long been known to produce IL8 upon stimulation with inflammatory mediators such as LPS (60). Increased levels of IL8 have been observed in secretions of male urethral challenge models of gonorrhea, which was postulated to be produced locally by the infected mucosa, rather than by infiltrating leukocytes (61). IPA enrichment of the Canonical Pathway, IL8 Signaling, was driven by the differential expression of CXCL1, CXCL8 (IL8), and VEGFA, all of which were observed to be induced by cytokine profiling (**Dataset S5, Fig 8A**). Since IL8 was near the upper limit of detection by cytokine array (**Fig 8A**), we measured concentrations of IL8 released from infected and uninfected PMNs by ELISA (**Fig 8B**). Here, IL8 pretreatment of PMNs was omitted to avoid confounding the detection of newly released IL8. Concentrations of IL8 released from PMNs significantly increased in response to infection with all three Gc strains but not in uninfected PMNs, beginning at 1 h and most pronounced at 2 h post-infection (**Fig 8B**). Thus, PMNs respond to Gc infection by releasing IL8, which is predicted to recruit and activate additional PMNs at the site of infection.

The Molecular Function Prostaglandin Synthesis (subset of Lipid Metabolism) was also significantly enriched, and PTGS2 (Prostaglandin-Endoperoxide Synthase 2, COX-2) was one of the most frequent genes driving Molecular Function enrichment (found in 11 of 15 functions) (**Figure 7, Dataset S5**). PTGS2 is induced during inflammation and synthesizes Prostaglandin H, which is highly unstable (62). Prostaglandin H is rapidly converted to Prostaglandin E2 (PGE2) by the constitutively expressed Prostaglandin E Synthase 2 (62), and PGE2 production switches neutrophils to an anti-inflammatory state to promote inflammation resolution (63). We measured the release of PGE2 from neutrophils exposed to Gc by ELISA (**Fig 8C**). Concentrations of PGE2 significantly increased following infection of neutrophils with all three Gc strains, but not uninfected neutrophils, beginning at 1 h and increasing at 2 h post-infection (**Fig 8C**). These findings suggest that production of PGE2 modulates the outcome of inflammation at sites of Gc infection, a hypothesis to be tested in future studies.

Upregulation of adhesion factors, particularly the integrins ICAM1 and ITGAX (CD11c), drove enrichment for multiple Molecular Functions and Canonical Pathways (**Fig 7, Dataset S5**). Interestingly, neither are considered conventional neutrophil proteins, yet both were transcribed at high levels (**Dataset S4**). ICAM1 is canonically expressed on endothelial cells to enable the extravasation of leukocytes from the blood into tissues but it has also been reported to be expressed by neutrophils, where it modulates effector functions such as ROS production and phagocytosis (64, 65). ITGAX (CD11c) is canonically considered a dendritic cell marker; however, neutrophils have recently been found to acquire antigen-presenting capabilities, including the induced expression of ITGAX, upon stimulation with select cytokines (66). We therefore assessed whether these were truly upregulated by neutrophils upon infection with Gc or were upregulated due to the presence of other cell types present at low levels in our purified PMN population. ICAM1 and CD11c protein expression on the surface of PMNs following exposure to all 3 strains of Gc at 1 and 2h post-infection was measured by flow cytometry (**Fig 8D and 8E**). Both ICAM1 and CD11c were expressed on purified human neutrophils (CD11b^+^/CD14^-^/CD16^+^/CD49^-^ cells) and surface levels of ICAM1 were significantly increased on neutrophils following Gc exposure (**Fig 8D**). In contrast, despite the increase in ITGAX transcripts measured in the Time-Adjusted PMN gene list, surface detection of CD11c was significantly decreased at 1 h post-infection (**Fig 8E**), suggesting that it may be internalized during infection with Gc, possibly during phagocytosis.

Overall, the Time-Adjusted PMN DE gene list indicates neutrophils have a concerted transcriptional response to Gc infection, resulting in upregulation of surface and secreted proteins that mediate neutrophil chemotaxis, phagocytosis, and resolution of inflammation. These analyses also reveal previously unknown pathways that are altered during Gc infection of neutrophils for future investigation.

## Discussion

Symptomatic gonorrhea is characterized by the influx of PMNs to the site of infection, but despite PMNs’ potent antimicrobial activities, viable Gc are recovered from PMN-rich exudates. The mechanisms behind the survival of Gc in this hostile environment have not been fully elucidated. Using dual RNA-seq profiling of Gc and human primary PMNs in a reductive model of acute infection, we performed a comprehensive analysis of the early shared transcriptional response across three unrelated individuals, and three isolates of Gc. We discovered that the Gc transcriptional response to PMNs overlaps with the transcriptional response to nutrient metals and oxidative stress, and this response impacts Gc survival during infection. Simultaneously, we identified host inflammatory and migratory responses as central to the host transcriptional response against Gc. These findings inform the complex ways that Gc and PMNs respond to one another during co-culture and reveal targets for future mechanistic studies and potential therapeutic intervention.

The transcriptomic analyses were conducted using two different strain backgrounds (**Fig S3**): FA1090 (isolated in the 1980s and used extensively in lab experimentation, the background for 130 and 3×130) and H041 (first reported in 2011 as a multi-drug-resistant isolate)(10, 23). PMNs responded similarly towards Gc regardless of strain background (**Fig 3B**). While we focused on analysis of the core Gc response to PMN exposure (**Fig 3A, PC1**), some differences between strain backgrounds were also observed (**Fig 3A, PC2**). These differences may be explained in part by the derepression of the *mtrR* regulon in H041 and 3×130 (**Dataset S3**), which harbor a mutation in the promoter for the *mtrR* repressor(10). Future studies will mine these data to examine the impact of antimicrobial resistance genes on strain-specific transcriptional responses during interactions with PMNs.

We found that Gc differentially expresses a variety of genes involved in metabolism and nutrient acquisition in response to infection of PMNs, in categories including nitrogen metabolism, ABC transporters, and metal acquisition. We anticipate these changes indicate, in part, the shift from growth in the nutrient rich medium GCBL (0h), to growth in a less rich medium for Gc, RPMI + 10% FBS (1 h). For this reason, we contextualized the 118 Gc genes identified as DE over time (Gc+PMN_1h vs Gc+PMN_0h) with those genes DE in the absence of PMNs (Gc+PMN_1h vs Gc_1h). This contextualization revealed Gc transcriptional responses that are specific to culture with PMNs that were otherwise masked by the shift in media. The transcriptional response reported here differed substantially from the response of Gc strain 1291 to PMNs in suspension, which showed only 33 genes identified as DE between 10 and 180 minutes of incubation with PMNs (32). Eleven of these 33 genes were DE in Gc upon exposure to adherent PMNs after 60 minutes in our analysis. This difference may reflect the enhanced ability of adherent PMNs to respond to Gc compared to PMNs in suspension (38), which are poorly activated and unable to internalize unopsonized, Opa^-^ Gc (67). Additional differences could be due to strain background, media, or timepoints.

Exposure of Gc to PMNs induced a prominent iron response, representing nearly 20% of all the DE genes identified. This, in part, represents iron restriction during ex vivo infection conditions, in which iron availability for Gc in the culture medium (RPMI+10% FBS) is limited due to the specificity of the Gc TbpAB for human transferrin over bovine transferrin (53, 54). Gc is capable of pirating iron from the human host from transferrin, lactoferrin, and hemoglobin in a TonB-dependent manner (83). We found that a *tonB* mutant had a survival defect in the presence of PMNs, implying that metal acquisition through TonB is required to resist PMN killing. While we cannot currently attribute the survival defect of Δ*tonB* solely to limitation of iron compared with other essential metals such as zinc, we also found that incubation in RPMI also induces a transcriptional response in Gc that is consistent with iron starvation (29). Intriguingly, the iron-starvation response is dampened when Gc is incubated with PMNs **(Fig. 5, Dataset S3)**. Of the 23 genes DE across Gc in response to PMNs, two were components of the TonB system, ExbB and ExbD, and at least four were iron-regulated components of TonB-dependent transporters (TdfF, TdfJ, TbpA, FbpA) (68). Together, these data suggest that iron limitation is a major mediator of Gc transcriptional responses towards PMNs, which induce expression of the TonB system and the transporters it energizes. Gc can acquire iron from human iron-binding proteins using TbpAB (transferrin binding protein AB), LbpAB (lactoferrin binding protein AB), and HpuAB (hemoglobin-haptoglobin utilization proteins) (68). Of these, TbpAB is likely to be the primary means of iron acquisition during PMN challenge, because LbpAB, although present and functional in H041, is truncated and nonfunctional in the FA1090 background (68), and erythrocytes are lysed and their contents separated from PMNs during purification, removing hemoglobin as an available iron source. While TbpB transcript and protein expression were highest in the inoculated Gc (Gc+PMN_0h), after 1 h there was more expression in unexposed Gc (Gc_1h) compared with Gc exposed to PMNs (Gc+PMN_1h). To our knowledge, this is the first evidence to suggest that Gc directly obtains iron from PMNs to overcome human nutritional immunity.

Despite Gc encoding numerous oxidative stress defenses (14), only four of the dedicated oxidative stress resistance genes in Gc were upregulated in response to PMNs: NGO2059 (*msrA*), NGO1767 (*kat*), NGO1769 (*ccpR*) and NGO0168 (*znuA*/*mntC*) (14). Instead, many DE genes have secondary roles in oxidative stress resistance and include metabolic transporters such as sulfite/cystine/cysteine transporters (NGO2011-2014, NGO0373-0374, NGO0455-0456), nitrogen metabolism, and metal transporters. We found that exposure to PMNs enhanced the resistance of Gc to H_2_O_2_. This response may reflect that Gc exposed to PMNs have an enhanced need for nutrients that resist and repair oxidative damage. Alternatively, Gc exposed to PMNs may sense that they are in a hostile host environment and respond by increasing the expression of this cohort of genes. Future studies with mutants in these transport/metabolism systems will help to discriminate between these possibilities.

We were interested to find that there is substantial overlap in the Gc transcriptional response to PMNs, iron limitation, and H_2_O_2_. As has been previously reported (31), we also noted substantial overlap between the H_2_O_2_, iron, and anaerobic transcriptomes. While iron limitation was expected since there is no exogenous source of metal that Gc can use in our infection conditions, the H_2_O_2_ response was less expected since PMNs do not release high levels of ROS in response to the Opa^-^ Gc used in this study (69, 70). Moreover, antioxidant gene products are dispensable for Gc to survive exposure to PMNs (41). Instead, our data indicate that Gc associated with PMNs have increased expression of genes involved in free radical scavenging, which implies Gc is confronted with free radicals during PMN exposure and has a transcriptional program to resist them.

Historically, human PMNs have been considered to be transcriptionally limited compared to other immune cell types (71, 72); thus, we hypothesized that any DE human transcripts were likely to be important in the context of Gc infection. Our results detail a response of PMNs to Gc that was independent of donor and bacterial strain background. We measured upregulation of chemotactic and immune-modulatory gene products in response to Gc, in particular, increased release of the proinflammatory cytokines CXCL-8 (IL8), IL1A, IL1B, IL1RA, MIP1A (CCL3), MIP1B (CCL4), and CXCL1 (GRO), and increased surface exposure of ICAM1. These factors are known to be associated with pro-inflammatory pathways (73) and confirms that the core response of PMNs to Gc includes an inflammatory component. Yet we also observed upregulation of PTGS2 and increased release of its product PGE2, which contribute to resolution of inflammation (63). These results underscore the complexity of the PMN immunomodulatory response to Gc infection.

Our findings align with a previous report from Sintsova *et al*. on the inflammatory response of PMNs to Gc (74). PMNs from CEACAM-humanized mice infected with Gc exhibited upregulation of proinflammatory pathways and genes, including TNFA, IL1A, CXCL1 (Gro-A/KC), MIP1A and MIP1B, PTGS2 (COX-2), NLRP3, and OSM (74). Production of pro-inflammatory cytokines by mouse PMNs was enhanced by Opa/CEACAM interactions (Opa^+^ Gc and CEACAM-transgenic PMNs), which was also confirmed with human PMNs infected with Opa^+^ vs. Opa^-^ Gc. These results suggest that the proinflammatory responses we observed with Opa^-^ Gc may be further enhanced during infection with Opa^+^ Gc, and may explain some of the differences observed between PMNs exposed to the ΔOpa 130 background compared to H041 which exhibits variable Opa expression.

IPA analysis was limited by a paucity of pathways directly associated with neutrophils. Gene set enrichment analysis (GSEA, **SI Appendix**) suffered the same limitation. Therefore, IPA analysis was set to include pathways attributed to all primary cell types, and we focused our analysis on Molecular Functions, which are more widespread between cell types, rather than Canonical Pathways, which are limited to specific cell types and disease states. Focusing our analysis on all primary cell types resulted in several genes repeatedly driving enrichment of many Molecular Functions, for instance, IL1B was involved in all of the top 15 Molecular Functions, marking these genes as highly interactive partners in the PMN inflammatory response during Gc infection. An additional strength of this approach is that previously unappreciated PMN functions can be revealed, for example implicating ICAM1 and ITGAX. One consequence of this approach, however, is that misleading pathways that are highly studied, but not relevant to PMNs, are enriched. For example, 8 different Canonical Pathways related to IL17 signaling were identified by IPA, however the production of any IL17 family members by neutrophils has recently been questioned (75). Further examination of our dataset revealed that, although IL17C was induced (L2FC of 2.75-3.65), normalized expression levels were exceptionally low for IL17C compared to genes for known cytokines produced by neutrophils i.e. CXCL8 (**Dataset S5**). Differential expression calculations for lowly expressed genes are unreliable, and should therefore be carefully validated. Our analysis focused on the top 20 most highly expressed PMN genes to account for this caveat, however the growing importance of PMNs in diverse inflammatory and infection conditions warrants reworking of available analytical pipelines to better assess rare transcripts for these critical cell types.

Interestingly, we did not observe substantial cross-talk between Gc and PMNs at the transcriptional level. For example, although we observed the upregulation of Gc metal acquisition genes, we did not identify corresponding host transcripts for the manipulation of metal trafficking in the host. Despite this lack of transcriptional upregulation, metal sequestering proteins like lactoferrin and calprotectin (S100A8/S100A9) are highly abundant as pre-formed proteins within the PMNs, and have well established functions in metal sequestration during Gc infection (52). This lack of cross-talk may reflect, in part, the abundance of preformed proteins in PMNs residing within cytotoxic granules that can rapidly carry out their effects without the need for transcriptional induction. Alternatively, the absence of cross-talk may indicate that PMNs have an initial somewhat non-specific response to Gc that involves the kinds of inflammatory pathways they would direct towards any other bacteria. The ability to circumvent both broad and specific PMN antimicrobial responses through unique transcriptional programs mark Gc as a successful pathogen.

Our analysis using multiple human donors and multiple isolates of Gc that differ in antimicrobial resistance status allowed us to identify a core response to infection in both Gc and in PMNs. The dual transcriptomes of Gc and human PMNs generated here provide an unbiased foundation for subsequent investigations into host-pathogen interactions during PMN challenge and will serve as a critical reference to support further human infection studies by microbial pathogenesis research communities. Our study highlights nutritional immunity and modulation of inflammation as key features of the Gc-PMN interface. These findings can reveal new bacterial and host targets for antimicrobial therapy, vaccine design, and prevention of inflammatory damage in the context of drug-resistant gonorrhea.

## MATERIALS AND METHODS

### Bacterial strains and growth conditions

Gc strain 130 is a non-variable Opa-deficient (Opa^-^) derivative of the FA1090 background constitutively expressing the pilin variant 1-81-S2 (23, 76). OpaD Gc is an isogenic derivative of 130, constitutively expressing OpaD (23). Strain H041 was received from R. Nicholas (UNC). Strain 3×130 was constructed by transformation of the *penA, mtrR*, and *penB* alleles from H041 into 130 (see Supplemental Methods). The pVCU915 plasmid containing the *tonB::kan* mutant from C. Cornelissen (53, 77) was transformed into 130. Gc were grown on Gonococcal Medium Base (GCB, Difco) plus Kellogg’s supplements (78) at 37°C with 5% CO_2_. For neutrophil experiments, Gc were grown in liquid medium (GCBL) for successive rounds of dilution to mid-logarithmic phase and enriched for piliation, as previously described (69). The genomes of 130, 3×130, and H041 used in this study were sequenced using PacBio technology (NCBI BioProject PRJNA508744, Genbank whole genome shotgun accessions WHPH00000000, WHPI00000000, and WHPL00000000).

### Gc-PMN co-culture

PMNs were isolated from venous blood as previously described (19) and used within 2h of isolation. Subjects gave informed consent in accordance with an approved protocol by the University of Virginia Institutional Review Board for Health Sciences Research (#13909). Synchronized Gc infection of adherent, IL8 treated PMNs was conducted as previously described (19). Gc-PMN samples were harvested immediately (Gc+PMN_0h) or after 1 h at 37°C with 5% CO_2_ (Gc+PMN_1h), by washing in ice-cold PBS and collecting in RNAprotect Cell Reagent (Qiagen) according to manufacturer instructions. Gc without PMNs at 1 h (Gc_1h) and PMNs without Gc (PMN_0h, PMN_1h) were also collected. Where indicated, CFU were enumerated from PMNs lysed in saponin at specified time points and expressed relative to the CFU at 0 h (100%).

### RNA extraction, library construction, and sequencing

RNA was extracted from the samples using the Qiagen RNA extraction kit with modifications, specifically the addition of 20 mM EDTA prior to lysis to inactivate PMN RNAses. Addition of EDTA reduced RNA degradation and improved RNA quality and quantity (**Dataset S1**). Bacterial and human rRNAs were depleted, 300 bp-insert strand-specific RNA-seq Illumina libraries were constructed, and RNA-seq was conducted on 150 nt pair-end runs using the Illumina HiSeq 4000 platform using two biological replicates for each condition. See **SI Appendix** for details, and **Dataset S1** for total number of reads generated per sample. RNA-seq data generated in this study are deposited at Gene Expression Omnibus (GEO) database under accession number GSE123434.

### Transcriptomic data processing

Illumina reads were trimmed for adaptor sequence and quality. Bacterial reads were mapped to the Opaless_3X (GenBank Accession: WHPL00000000) or H041 (GenBank Accession: WHPH00000000) genomes using bowtie v1.0 (79), and human reads were mapped to the 2018 GRCh38 Human genome assembly using HISAT v2.0 (80). For details on RNA-seq data analyses and tools used to generate Figures 2, 3, and 4, see **SI Appendix**, and **Data and Code Availability** below. All predictions of differentially expressed (DE) genes were performed using DEseq2 (81) and filtered using an FDR cutoff of 0.05 and an absolute Log_2_(Fold Change) cutoff of 1. Individual heatmaps of DE genes are available in **Dataset S2**, and gonococcal and human DE genes are listed in **Datasets S3** and **S4**, respectively.

Bacterial DE gene estimation was performed by comparing Gc+PMN_1h to Gc+PMN_0h and to Gc_1h (See **Figure 4**). For the Gc+PMN_1h vs Gc+PMN_0h comparisons, genes that were present in both the FA1090 background strains and H041 (core genes, **Dataset S3 Tab 7**) were considered. Core DE genes were subjected to KEGG pathway analysis (**Table S2**, see **Data and Code Availability**) as well as a manual literature survey.

For PMNs, genes that were DE in Gc+PMN_1h vs Gc+PMN_0h for each of the three Gc strains, and in PMN_1h vs PMN_0h, were analyzed. Genes that were DE in PMN_1h vs PMN_0h were not included, in order to focus specifically on genes DE in response to Gc. To capture genes with magnitude differences during infection but also changing with time, 632 time-dependent genes (common to all four 1 h vs 0 h comparisons, **Fig 6A - purple bar**) were intersected with DE genes in Gc+PMN_1h vs PMN_1h comparisons (**Fig 6B**). The resulting 106 genes (**Fig 6B - green bar**) were added to the 303 genes that were DE for PMNs infected with Gc over 1 h (Gc+PMN_1h vs Gc+PMN_0h), yielding the Time-Adjusted PMN list of 409 DE genes **(Fig 1A and 1B – sum of green bars, Dataset S4, Fig S5**). Gene identifiers as well as average Log_2_(Fold Change) were used as input for Ingenuity Pathway Analysis (IPA) (Qiagen, Redwood City CA) where IPA-defined core analyses for Canonical Pathways and Disease and Biological functions were performed using all primary cell subsets (**Figure 1)** (82) (QIAGEN Inc., https://digitalinsights.qiagen.com/IPA). The Time-Adjusted genes were also subjected to Gene Set Enrichment Analysis (GSEA) (83, 84) (see **SI Appendix**).

### Quantification of Cytokine Secretion

Multiplexed cytokine quantification was profiled using the 38-plex Human Cytokine/Chemokine Magnetic Bead Panel on the Luminex MAGPIX instrument (Millipore). IL8 was measured with Human IL8/CXCL8 DuoSet ELISA (R&D Biosystems) in PMN supernatants diluted 1:4. PGE2 was measured by Prostaglandin E2 (Highly Sensitive) ELISA (Immuno-Biological Laboratories, Inc.) using undiluted samples.

### Flow cytometry

Surface expression of ICAM1 (CD54) and ITGAX (CD11c) was measured on PMNs (CD49d+/CD16+/CD11b+/CD14-) by flow cytometry using Cytek Northern Lights spectral flow cytometer, and data were analyzed with FCS Express (De Novo Software, Pasadena, CA). Fluorescence minus one (FMO) controls were used to set gates for analysis.

### H_2_O_2_ sensitivity assays

130 Gc was exposed to adherent, IL8 treated PMNs for 1 h as described above. PMNs were lysed in 1% saponin, and Gc were pelleted, washed, and resuspended in GCBL at 10^7^ CFU/ml to yield the pre-exposed bacterial population. The concentration of Gc was based on the expected recovery of viable bacteria (**Fig. S1**) and was verified by CFU enumeration. Pre-exposed and naïve (GCBL-grown) Gc were exposed to the indicated concentrations of H_2_O_2_ (Sigma) in GCBL and incubated with rotation for 15 minutes at 37 ºC. Bacteria were then immediately diluted into GCBL containing bovine catalase (final concentration of 10 μg/ml; Sigma) to degrade residual H_2_O_2_ and plated at limiting dilution on GCB agar. CFU/mL was derived from enumerated colonies, and percent viability was calculated by dividing CFU/mL at each time point by CFU/mL at 0 min. Significance determined by linear mixed effects model: Survival ∼ Treatment + (1 | Subject) + H_2_O_2_ Concentration.

### TbpB expression

Gc were exposed to adherent, IL8 treated PMNs for 1 h as described above. PMNs were lysed in 1% saponin, and Gc cells were pelleted (5 mins, 3000xg), washed in GCBL, and resuspended in 1X Laemmli sample buffer containing SDS and β-mercaptoethanol. Gc lysates were separated by 10% SDS-PAGE, transferred to nitrocellulose, and blotted for TbpB and Zwf as loading control (85) (86). Quantification of band intensity using infrared secondary antibodies and normalization was performed in ImageStudio Ver 5.2 (LI-COR) (see **SI Appendix**).

## Data & Code availability

R code/packages/scripts used to perform transcriptomics data analyses and generate figures are available on GitHub at https://github.com/admelloGithub/PMN_GC. All RNA-seq data have been deposited in the Gene Expression Omnibus (GEO) database under accession GSE123434.

## Author contributions

A.K.C. and H.T. designed research; V.L.E., A.D.P., A.D., M.G., X.Z., A.K.C., and H.T. performed research; K.H. and S.R. contributed new reagents; V.L.E., A.D.P., A.D., A.C.S., A.K.C., and H.T. analyzed data; V.L.E., A.D.P., A.D., A.K.C., and H.T. wrote the paper.

## Acknowledgments

We thank Jacques Ravel and Patrik Bavoil for initial discussions through the Ecopathogenomics of Sexually Transmitted Infections project (NIHU19AI094044) and David Rasko for enabling supplements to the Genomic Centers for Infectious Diseases project (NIHU19AI110820). We thank Robert Nicholas for the H041 strain and advice on construction of 3×130; Jane Wilhelm for critical help with RNA purification; Cynthia Cornelissen and Aleksandra Sikora for antibodies and mutant strains; Michael Solga, Taylor Harper, and Claude Chew (UVA Flow Cytometry Core) for advice; Marieke Jones and David Martin (UVA Claude Moore Health Sciences Library) for statistical analyses; and the Institute for Genome Sciences’ Genomics Resource Center, Genome Informatics Core, and High-Performance Computing Core for essential support. This project was funded in part by the National Institute of Allergy and Infectious Diseases, National Institutes of Health (NIH), Department of Health and Human Services under grant number U19AI110820. Additional funding support was from: NIH: R21AI130646 to V.L.E. and H.T.; T32AI007496 to A.D.P.; R01AI097312 and R21AI161302 to A.K.C.; U19AI094044 to A.K.C. and H.T.; UVA: Hutcheson Family Fellowship and Institute for Practical Ethics in Public Life Fellowship to K.H., and Robert R. Wagner Fellowship to S.A.R.

