## Supplemental Information Appendix for "Dual species transcriptomics reveals core metabolic and immunologic processes in the interaction between primary human neutrophils and *Neisseria gonorrhoeae* strains"

**This PDF file includes:**

Supplementary text
Figures S1 to S7
Tables S1 to S2
Legends for Datasets S1 to S6
SI References

**Other supplementary materials for this manuscript include the following:**

Datasets S1 to S6

### SUPPLEMENTAL TEXT

**GSEA ANALYSIS.** In addition to IPA analysis of pathways enriched in the 409 Time-Adjusted PMN DE genes, we subjected the same list of genes to Gene Set Enrichment Analysis (GSEA) (**Dataset S6**) (1). Very few datasets included in GSEA were associated with PMN or antibacterial responses, and a majority of the datasets deemed enriched by GSEA were cancer related. We did observe some overlap of Time-Adjusted PMN DE genes with those of inflammatory responses in other immune cell types by GSEA. Of the immune response associated datasets identified, most used microarrays (9-11) and their focus was mainly on cells exposed to various immune modulators or inhibitors (10-14). Despite these caveats, GSEA analysis confirmed that the main core functions enriched in PMNs exposed to Gc were signaling, immune response, cell cycle, and cell morphology. These results further validated the pathways observed in IPA. The enriched gene sets that were relevant to these responses are discussed here.

Gc produces lipooligosacharide (LOS), a derivative of lipopolysaccharide (LPS), as a major component of its outer membrane that is highly inflammatory. As such, we examined GSEA datasets associated with host responses to LPS. Human macrophages, upon exposure to live *P. gingivalis*, upregulated 458 genes, 52 of which overlap with Time-Adjusted genes (11.5% of that dataset) (ZHOU_INFLAMMATORY_RESPONSE_LIVE _UP) (3). Whereas exposure to purified *P. gingivalis* LPS resulted in upregulation of 409 genes in macrophages, 26 (6.4% of that dataset) overlapped with Time-Adjusted PMN genes (ZHOU_INFLAMMATORY _RESPONSE_LPS_UP) (3). Another dataset analyzed activation of myeloid derived dendritic cells (MDDCs) by Galectin-1 (known to play a role in modulation of immune cells) compared to LPS (FULCHER _INFLAMMATORY_RESPONSE_LECTIN_VS_LPS_UP) (2). Galectin-1 activated MDDCs had an overlap of 33 genes from the 409 Time-Adjusted PMN genes, and these 33 genes represent 5.6% of the 585 total genes activated in Galectin-1 treated MDDCs. In examination of dendritic cell maturation in which MDDCs were stimulated with monocyte conditioned media containing IL1B at early (8h) compared to late timepoints (48 h) (6), 16 genes (26.2%) that were upregulated at 8 hours overlapped with our dataset. A dataset examining neutrophils that had transmigrated towards a skin wound (THEILGAARD_NEUTROPHIL_AT_SKIN_WOUND _UP), overlapped with 17 (22.4%) genes (5).

Most relevant, in addition to experimental datasets, GSEA also performs enrichment and analysis for pathways in the reactome database. The degranulation of neutrophils reactome (REACTOME_NEUTROPHIL

_DEGRANULATION) (4) overlapped with 13 (2.7%) Time Adjusted PMN DE genes. Additionally, cytokine signaling (REACTOME_CYTOKINE_SIGNALING_IN_IMMUNE_SYSTEM) and interleukin signaling (REACTOME_SIGNALING_BY_INTERLEUKINS) had an overlap of 4.2% and 5.4% of genes observed.

### MATERIALS AND METHODS

**Construction of strain 3X130**. To construct 3X130, the *penA* allele from H041 was amplified by PCR using PenA F 5'-CAATCCGCATCTTTATTAACCGCGAGC-3' and PenA R 5'-CCGGTTTCAGCCAAAGGGCTTAAC-3'. The resulting product was transformed into 130, and transformants were selected with 40 ng/ml cefixime. The *mtrR* allele from H041 was amplified using mtrR F 5'-ACGGGTTGCAAAGCAGGTTATACC-3' and mtrR R 5'-CCCTTTCAAACGGCATCAAAATGACAC-3'. The resulting product was transformed into penA-130, and transformants were selected on 250 ng/ml erythromycin. The *penB* allele from H041 was transformed into penA/mtrR-130 by gDNA transformation, and transformants were selected on 20 ng/ml penicillin. The final resultant penA/mtrR/penB-130, termed 3X130, was confirmed by DNA sequencing of all three alleles (and subsequently by whole genome sequencing in this study). Antibiotic resistance was verified by E-Test strip compared to H041 and 130 (**Table S1**).

**RNA extraction from PMNs and Gc**. Adherent PMNs ($3.6*10^6$ PMNs/well) were treated with 10 nM human IL-8 (R&D Biosystems) in Roswell Park Memorial Institute 1640 medium (RPMI) with 10% fetal bovine serum (FBS) at 37°C with 5% $CO_2$ for at least 30 min prior to infection. Gc at an MOI of 5 were centrifuged onto PMNs chilled on ice to synchronize infection. Samples were collected in RNAprotect Cell Reagent (Qiagen) at the University of Virginia and shipped on dry ice to the University of Maryland.

After thawing on ice, 20 mM EDTA was added to each sample and incubated at room temperature for 15 min to inactivate RNAses released from neutrophils. Samples were pelleted at 13,000 x g for 5 min at 4°C, the supernatant removed, and 100 µl of lysis solution added (2.50 U/µl mutanolysin, 4 µg/µl Proteinase K, 3 µg/µl lysozyme, TE buffer 40 µl) plus 4 µl EDTA (500mM). These samples were vortexed and then incubated for 10 min at room temperature with intermittent vortexing every 2 min. Samples were extracted using the Qiagen RNA extraction kit as per manufacturer's recommendations with slight modifications. Briefly, 700 µl RLT buffer containing BME (10µl /ml) was added to each sample and vortexed for 30 sec. This was followed by centrifugation for 30 sec at 13,200 x g and the supernatant transferred to a new tube. 800 µl 70% ethanol (1 volume based on supernatant volume) was added and 700 µl transferred to a RNeasy spin column, spun at 8000 x g for 30 sec and flow through discarded. This step was repeated until all the sample passed through the column. 350 µl RW1 buffer was then added to the column and spun at 8000 x g for 30 sec, then 80 µl of

DNase solution (70 µl RDD buffer, 10 µl DNAse) added to each filter and incubated at room temperature for 15 min. 350 µl RW1 buffer with 2 0mM EDTA was added and incubated for 10 min at room temperature, spun at 8000 x g for 30 sec and the flowthrough discarded. 500 µl RPE was added to the column, incubated at room temperature for 5 min, spun at 8000 x g for 1 min and flowthrough discarded. The column was then transferred to a new collection tube, 50 µl Rnase-DNase free water added and incubated at room temperature for 5 min. followed by a spin at 8000 x g for 1 min and the RNA collected.

To remove any residual DNA, and additional Dnase treatment was conducted in solution. 5 µl 10X Turbo DNAse buffer and 1 µl DNA free enzyme (2U/µl) were added to the sample and incubated at 37°C for 30 min. An additional 1 µl DNA free enzyme (2U/µl) was added and samples incubated for an additional 30 min at 37 °C. 12 µl DNAse inactivation reagent was added and tube flicked for even distribution. Samples were incubated for 5 min at room temperature with intermittent flicking and centrifuged at 10,000 x g for 2 min. Supernatant (RNA) was transferred to a new tube and stored at -80 °C.

**RNA quantification.** 1 µl of sample or RNA ladder was added to 5 µl RNA Tape Station buffer (Agilent; with a maximum buffer concentration of 200 mM Tris, 20 mM EDTA, 50 mM NaCl per sample) as per manufacturer's instructions. Samples were heated to 72°C for 3 min, followed by incubation on ice for 2 min. Samples were analyzed on the Tape Station instrument as per Agilent recommendations and RNA Integrity Numbers (RIN) and RNA concentrations determined (**Dataset S1**).

**Acquisition of high-quality RNA from PMNs.** Extraction of high quality and quantity RNA from PMNs is not trivial, as neutrophil granules contain RNases that are released upon cell lysis (15). The standard metric of RNA quality for sequencing (RNA Integrity Number or RIN) is recommended to be over 8 for high quality RNA samples. Conventional RNA isolation from samples containing PMNs gave low quality RNA, with RIN scores as low as 3.6 with a concentrations as low as 1.6 ng/µl. We found that addition of EDTA prior to cell lysis reduced RNA degradation and improved RNA quality and quantity, resulting in samples with average RIN scores of 5.38 for samples containing PMNs alone, 5.0 for samples containing PMNs and Gc, and 8.9 for samples containing Gc alone (**Dataset S1**). RNA concentrations were also improved with EDTA treatment, with Gc alone samples, averaging 77.4 ng/µl, while those containing PMNs averaged 26.8 ng/µl (**Dataset S1**).

The use of EDTA treatment allowed for RNA of sufficient quality and quantity to be collected for the preparation of libraries and subsequent sequencing.

**RNA library construction and sequencing.** 300 bp-insert strand-specific RNA-seq Illumina libraries were constructed using RNA that was enriched for mRNA by depletion of ribosomal RNA using the Ribo-Zero rRNA Removal Kits for Gram-negative bacteria and/or for human/mouse/rat (Illumina). RNA that was then fragmented and used for synthesis of strand-specific cDNA using the NEBnext Ultra Directional RNA Library Prep Kit (NEB-E7420L). The cDNA was purified between enzymatic reactions and the size selection of the library performed with AMPure SpriSelect Beads (Beckman Coulter Genomics). The titer and size of the libraries was assessed on the LabChip GX (Perkin Elmer) and with the Library Quantification Kit (Kapa Biosciences). RNA-seq was conducted on 150 nt pair-end runs of the Illumina HiSeq 4000 platform using two biological replicates for each condition.

**Transcriptomic data processing.** *Neisseria gonorrhoeae*: Sequence reads were trimmed for adaptor sequence and mapped to the Opaless_3X (GenBank Accession: WHPL00000000) or H041 (GenBank Accession: WHPH00000000) Neisseria genomes using bowtie v1.0 (16) with parameters -mode=v --num_mismatches=2 --file-type=fastq --seedlen=28 --minins=0 --maxins=600 --library-type=fr --args='--sam' --v --gzip .

*Homo sapiens*: Sequence reads were trimmed for adaptor sequence and mapped to the 2018 GRCh38 Human genome assembly using HISAT v2.0 (17) with parameters -mismatch-penalties=6,2 --softclip-penalties=2,1 --read-gap-penalties=5,3 --ref-gap-penalties=5,3 --min-intronlen=20 --max-intronlen=500000 --score-min=L,0,-0.2 --pen-cansplice=0 --num-threads=1 --pen-noncansplice=12 --pen-canintronlen=G,-8,1 --pen-noncanintronlen=G,-8,1 --rna-strandness=RF --dta-cufflinks=1 --num-alignments=5 --minins=0 --maxins=600 --no-unal=1 --args='' --v .

Normalized gene expression values and differential gene expression analyses for host and bacterial gene features were calculated as previously described (18). Briefly, gene expression counts for all samples were estimated using HTseq (19). For bacterial genes, counts were then tabularized and counts tables were imported into Rstudio for calculation of differentially expressed (DE) genes (see **Data and Code Availability** in the main document). Rarefaction curves in Figure 2 were generated from HTseq counts data (see **Data and Code Availability**). Principal Component Analyses (PCAs) and dendrograms in Figure 3 and Supplemental Figure 4 were generated using R studio based on normalized Variance Stabilized Transformation (VST) counts acquired using the DESeq2 R package (20). VST counts are transformed gene counts data, that have been normalized for sequencing depth and variance across biological replicates. All predictions of differentially expressed (DE) genes were performed using DEseq2 and filtered using an FDR cutoff of 0.05 and an absolute $Log_2$(Fold Change) cutoff of 1. DE genes shared between different comparisons or specific to a given comparison were determined using Upset plots (See **Data and Code Availability**). Individual heatmaps of DE genes were generated based on Z-scores of average VST counts per condition (**Dataset S2**). Gonococcal and human DE genes are listed in **Datasets S3** and **S4**, respectively.

**Gc analysis.** DE genes were first estimated for the comparisons of 130+PMN_1h vs 130_1h, 130+PMN_1h vs 130+PMN_0h, 3X130+PMN_1h vs 3X130_1h, 3X130+PMN_1h vs 3X130+PMN_0h, H041+PMN_1h vs H041_1h, and H041+PMN_1h vs H041+PMN_0h using all 3X130 or H041 genes (**Dataset S3, Tabs 1-6**). For all Gc+PMN_1h vs Gc+PMN_0h conditions, genes that were DE in both 3X130 and H041, and were also present in FA1090 (core genes, **Dataset S1 Tab 5**), were determined for each comparison. This revealed 23 core genes that were DE in all strains (**Fig 4, Dataset S3 Tab 8**), and 118 core genes DE in at least one strain (**Dataset S3 Tab 7**). The 118 genes were then subjected to KEGG pathway analysis (**Table S2,** see **Data and Code Availability**) as well as a manual literature survey (21-26), to reveal differentially regulated gonococcal pathways.

**PMN analysis.** DE genes from 4 comparisons – 130+PMN_1h vs 130+PMN_0h, 3X130+PMN_1h vs 3X130+PMN_0h, H041+PMN_1h vs H041+PMN_0h, and PMN_1h_vs_PMN_0h – were calculated. Genes that were DE in PMN_1h vs PMN_0h were not included, in order to focus specifically on genes DE in response to Gc. To capture genes with magnitude differences during infection but also changing with time, 632 time-dependent genes (common to all four 1h vs 0h comparisons, **Fig 6A - purple bar**) were intersected with DE genes in Gc+PMN_1h vs PMN_1h comparisons (**Fig 6B**). The resulting 106 genes (**Fig 6B - green bar**) were added to the 303 genes that were DE for PMNs infected with Gc over 1h (Gc+PMN_1h vs Gc+PMN_0h), yielding the Time-Adjusted PMN list of 409 DE genes (**Fig 1A and 1B – sum of green bars, Dataset S4, Sup Fig 5**). Gene identifiers as well as average $Log_2$(Fold Change) were used as input for Ingenuity Pathway Analysis (IPA) (Qiagen, Redwood City CA) where IPA-defined core analyses for Canonical Pathways and Disease and Biological functions were performed using all primary cell subsets (**Figure 1**) (27) (QIAGEN Inc., https://digitalinsights.qiagen.com/IPA, Dec 2021 release). Gene identifiers were linked to their appropriate gene object in the IPA knowledge base and the following specific parameters used: Ingenuity knowledge base genes only; Interactions include endogenous chemicals; All node types; All data sources; Confidence experimentally observed; Species human; Tissues and human cells (excluded cell lines); All mutations. To confirm that the probability that each biological function, disease or pathway predicted to be involved was not due to chance alone a Fisher's exact test was performed. IPA then computed a score for each network based on the set of genes input which indicate activation or inhibition of the function or pathway (27). From these data, we analyzed in detail both IPA core analyses for Canonical Pathways and for Molecular Functions categories within Disease and Biological functions with a cutoff of -log(pvalue 1.3) that corresponds to a pvalue <0.05 (**Figure 1**).

**GSEA Analysis.** The 409 Time-Adjusted PMN DE genes were subjected to GSEA v4.1.0, using the C2.all.v7.4.symbols.gmt(curated) database and default parameters: 1000 permutations, genes were collapsed, weighted enrichment, and signal2noise as the metric for ranking the genes. We specified that gene sets smaller than 5 or larger than 200 be excluded.

**Multiplexed cytokine detection.** Adherent PMNs were incubated with Gc as described above or left uninfected. PMN supernatants were harvested after 4h of incubation at 37°C with 5% $CO_2$, then stored at -20°C. PMNs were not pretreated with IL-8 to avoid confounding exogenously added IL-8 and IL-8 released in response to Gc; no differences were noted in the ability of PMNs to adhere or interact with Gc. Quantification of secreted cytokines was done by Millipore 38-Plex Human Cytokine/Chemokine Magnetic Bead Panel (HCYTMAG60PMX38BK) on the Luminex MAGPIX instrument and reported as pg/mL.

**Enzyme linked immunosorbent assays.** Adherent PMNs were incubated with Gc as described above or left uninfected. PMN supernatants were harvested sequentially at baseline (0h) or after 1 or 2h of incubation at 37°C with 5% $CO_2$, then stored at -80°C. For the IL-8 ELISA, PMNs were not pretreated with IL-8 to avoid confounding exogenously added IL-8 and the IL-8 released in response to Gc; no differences were noted in the ability of PMNs to adhere or interact with Gc. IL-8 was measured by Human IL-8/CXCL8 DuoSet ELISA (R&D Biosystems) with supernatants diluted 1:4. PGE2 was measured by Prostaglandin E2 (Highly Sensitive) ELISA (Immuno-Biological Laboratories, Inc.) using undiluted samples.

**Flow cytometry.** Adherent, IL-8 treated PMNs were incubated with Gc or left uninfected. PMNs were lifted with 5mM EDTA at 0, 1, and 2h post-incubation. Approximately $1x10^6$ PMNs were washed and stained with with PE/Cyanine5 anti-human CD11b Antibody Clone ICRF44 (Biolegend, San Diego, CA) and BV711 Mouse Anti-Human CD14 Clone MφP9 (BD Horizon) to identify polymorphonuclear cells (PMNs) from monocytes, and Brilliant Violet 421™ anti-human CD16 Antibody clone 3G8 (Biolegend, San Diego, CA) and PE/Cyanine7 anti-human CD49d Antibody Clone 9F10 (Biolegend, San Diego, CA) to discriminate neutrophils from eosinophils, by flow cytometry. ICAM1 and ITGAX (CD11c) surface expression was assessed by Median Fluorescent Intensity (MFI) of PE anti-human CD54 Antibody clone HCD54 (Biolegend, San Diego, CA) and APC anti-human CD11c Antibody clone 3.9 (Biolegend, San Diego, CA) on $CD11b^+/CD14^-/CD16^+/CD49^-$ cells, i.e. neutrophils. Antibodies were diluted in Brilliant Stain Buffer (BD Horizon) and incubated with samples on ice for 30 min. Samples were also stained with isotype control antibodies to detect non-specific binding. Samples were then resuspended in 2% paraformaldehyde (PFA) until processing by flow cytometry on the Cytek Northern Lights spectral flow cytometer in the UVA Flow Cytometry Core Facility. Data were analyzed using FCS Express (De Novo Software, Pasadena, CA). Fluorescence minus one (FMO) controls were used to set gates for analysis.

**Western blotting.** 130 Gc was exposed to adherent, IL-8 treated PMNs for 1h as described above. PMNs were lysed in 1% saponin, and Gc cells were pelleted (5 mins, 3000xg), washed in GCBL, and resuspended in 1X Laemmli sample buffer containing SDS and β-mercaptoethanol. Samples were boiled for 5 minutes and sheared through a 25G needle, then stored at -80ºC. Gc lysates were separated by 10% polyacrylamide SDS-

PAGE and transferred onto nitrocellulose. Blots were blocked for 16h in Tris-buffered saline containing 0.05% Tween-20 and 5% nonfat dry milk. TbpB was detected using polyclonal rabbit antisera (a gift of C. Cornelissen, Georgia State University) as described (28), followed by goat anti-rabbit IgG coupled AlexaFluor 680 (Invitrogen). As a loading control, blots were probed with rabbit anti-Zwf (a gift of Aleksandra Sikora, Oregon State University) (29). Quantification of band intensity and normalization was performed in ImageStudio v5.2 (LI-COR).

**SUPPLEMENTAL FIGURES**

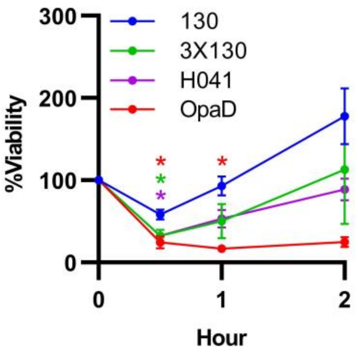


<u>Supplemental Figure 1:</u> **Gc experiences an outgrowth period at 1 hour following exposure to PMNs.** FA1090 Gc variants 130, 3X130, and OpaD, and Gc strain H041, were exposed to adherent, IL-8-treated primary human PMNs. Percent Gc survival was calculated by enumerating colony-forming units (CFU) from PMN lysates at 30, 60, and 120 min and reported as the percent of CFU for that strain at 0 min. Significance was determined by two-way ANOVA with Holm–Sidak correction for multiple comparisons. *indicates $p < 0.05$ compared to 130. $n = 4$ independent experiments.

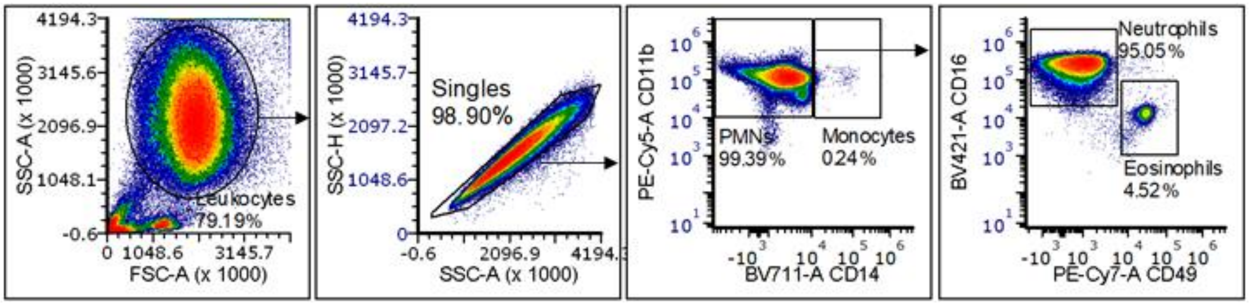


**<u>Supplemental Figure 2:</u> Purity of neutrophils in the human PMN preparation by flow cytometry**
IL-8-treated, adherent human PMNs were left untreated, or exposed to 130, 3x130, or H041 for 0, 1, or 2h. Cells were subsequently examined by flow cytometry using forward scatter (FSC-A) and side scatter (SSC-A) to identify single leukocytes, by PE-Cy5-CD11b and BV711-CD14 to distinguish polymorphonuclear cells (PMNs) from monocytes, and by BV421-CD16 and PE-Cy7-CD49 to distinguish neutrophils from eosinophils by flow cytometry. PE-ICAM1 and APC-CD11c was also assessed (Fig 8). Cytometry plots from one representative preparation of uninfected PMNs at 0h are shown.

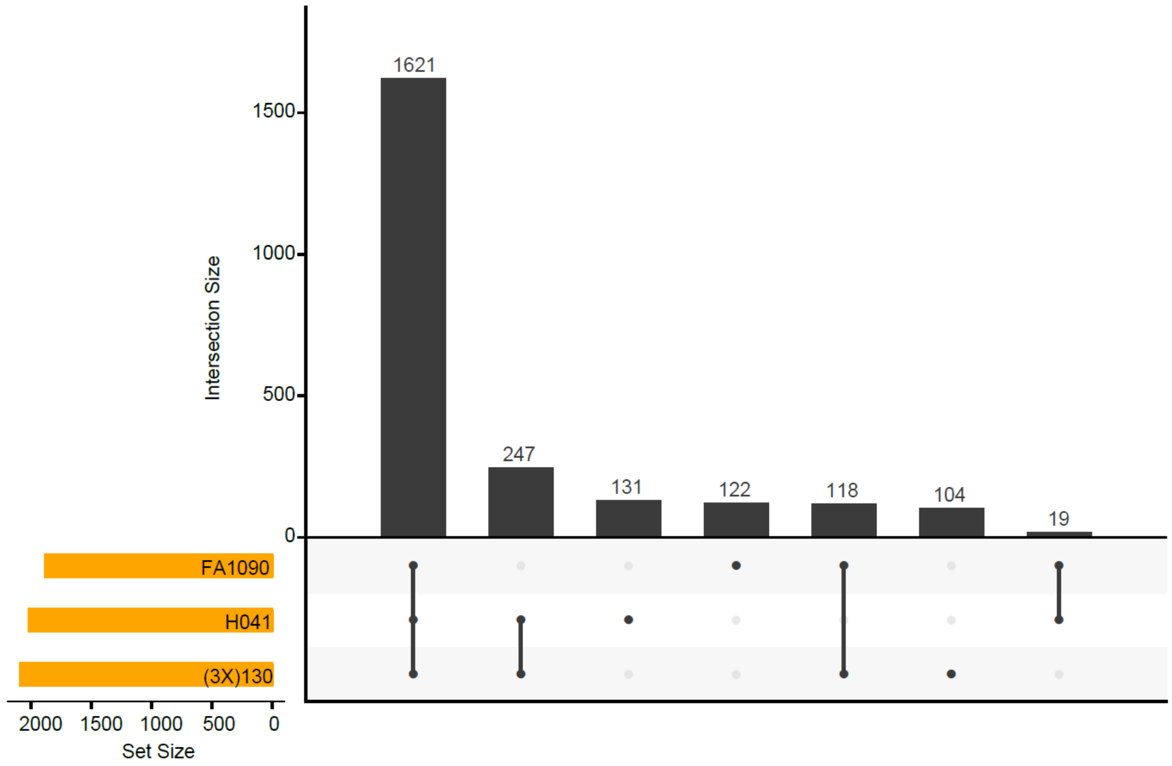


**<u>Supplemental Figure 3: </u>Identification of Gc core genes.** Orthologous (core) genes shared between strains 130, 3X130, H041, as well as the reference FA1090 genome sequence (Accession No. NC_002946.2, used for annotation purposes) were identified using PanOCT. Of the 1621 clusters of shared genes, clusters harboring paralogs (multiple genes from the same genome) and/or fragmented genes were removed, resulting in 1576 clusters of core genes used for Gc transcriptome analyses.

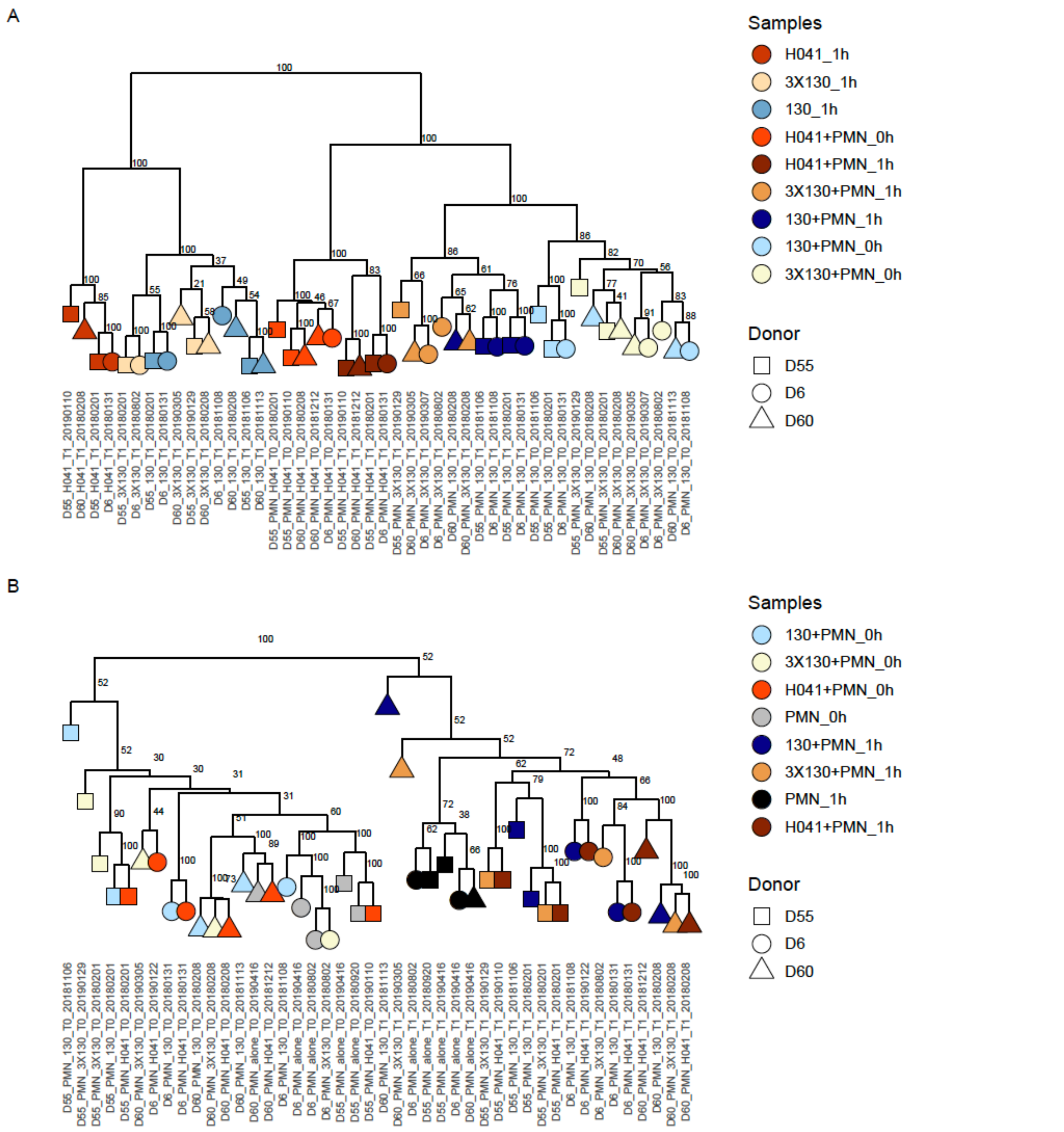


<u>**Supplemental Figure 4:**</u> **Gc and PMN dendrograms of expression profiles.** A) Bootstrapped dendrogram of Gc core gene expression profiles showing replicate clustering of samples by treatment condition. B) Bootstrapped dendrogram of PMN gene-expression profiles showing replicate clustering of samples by timepoint. Shapes are PMNs from different donors, colors are Gc strain or uninfected condition at the indicated timepoint.

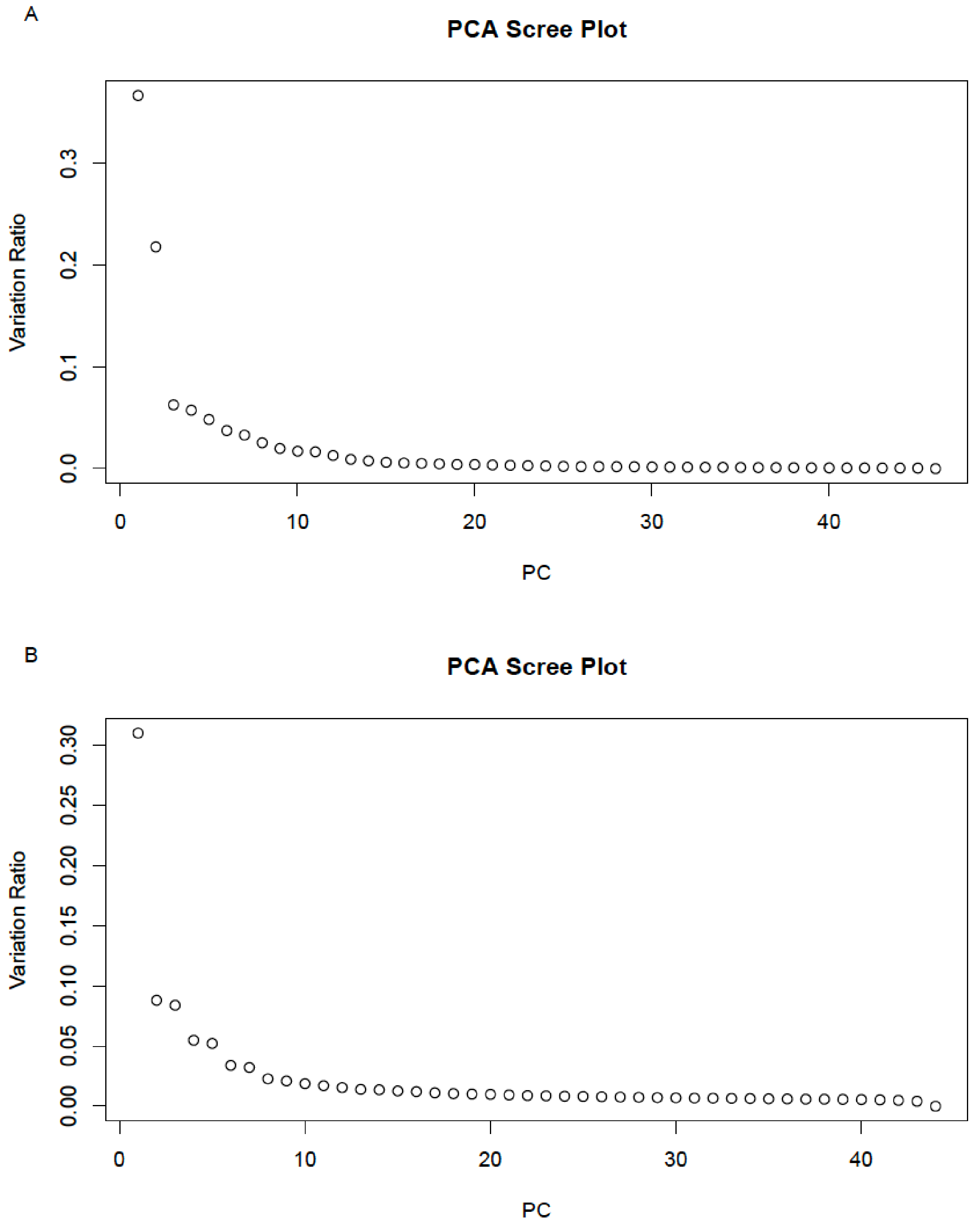


**<u>Supplemental Figure 5:</u> PCA Scree Plots.** A) PCA scree plot of Gc core expression profiles. The majority of Gc variability is captured by PC1 and PC2. Less than 10% of the Gc core gene expression profile is dictated by PC3 or above. B) PCA scree plot of PMN expression profiles. The majority of PMN variability is captured by PC1 and PC2 (Figure 3). Less than 10% of PMN gene expression profile is dictated by PC3 or above.

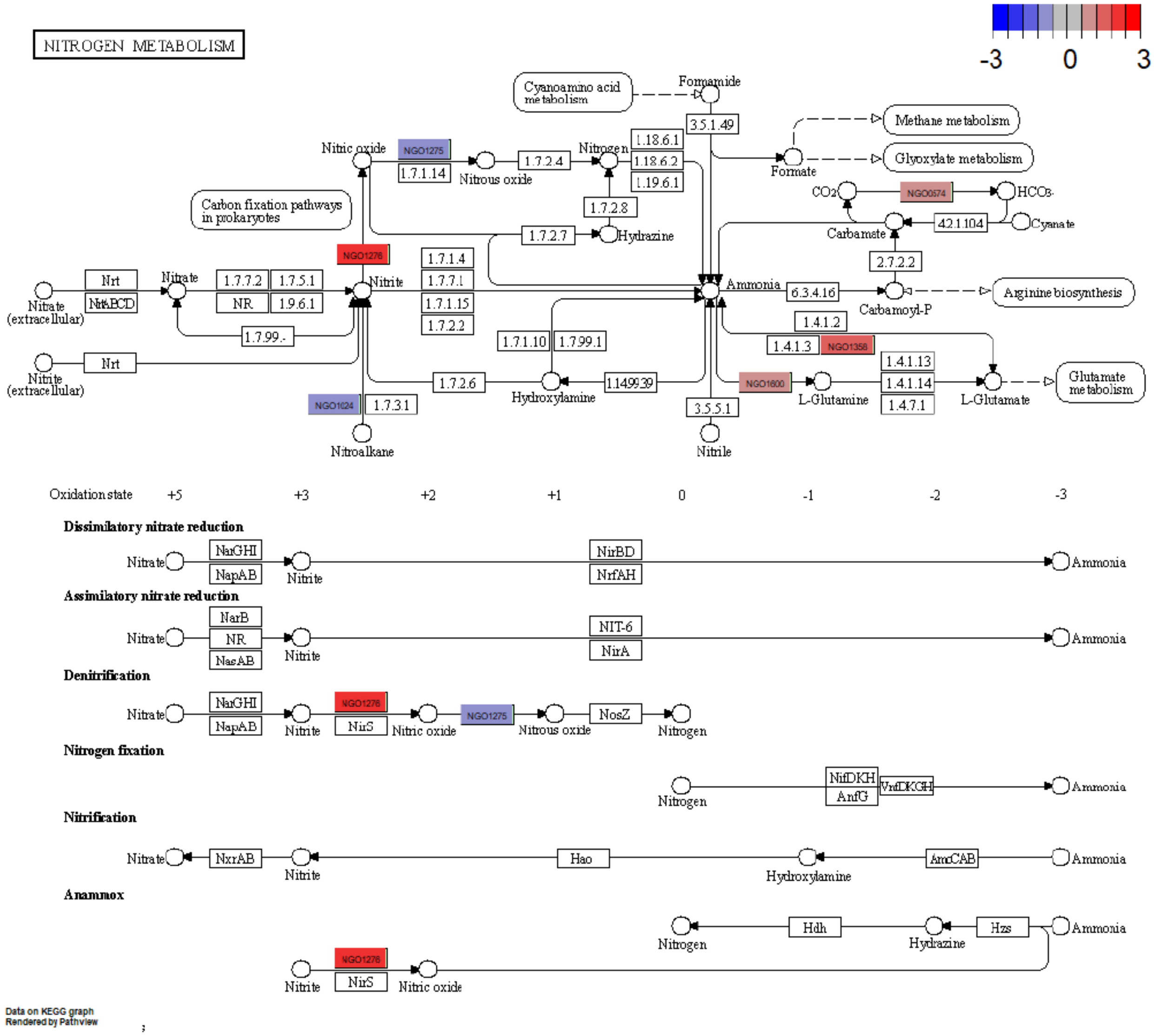


**<u>Supplemental Figure 6</u>: KEGG Pathway of Gc nitrogen metabolism.** Genes that were DE in Gc+PMN_1h vs Gc+PMN_0h were overlaid on the KEGG pathway for bacterial nitrogen metabolism. Colors indicate the respective average $Log_2$(Fold Change) for Gc genes, red is upregulated, blue is downregulated, white is N/A.

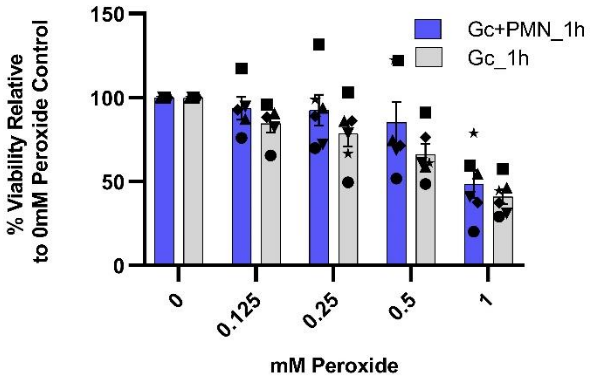


**Supplemental Figure 7: Gc resistance to hydrogen peroxide is enhanced by exposure to PMNs.** Gc strain 130 was inoculated onto IL-8-treated, adherent human PMNs or a media alone control, and incubated for 1 h. PMNs were treated with 1% saponin to liberate intracellular bacteria for 10 min before harvesting. Bacteria were washed and resuspended in GCBL at $10^7$ CFU/ml. Gc recovered from the PMNs (Gc+PMN_1h) or from growth in media alone (Gc_1h) were then exposed to 1, 0.5, 0.25, 0.125 or 0 mM $H_2O_2$ for 15 min, before quenching with catalase and plating at limiting dilution for CFUs. Shapes are data points from different donors' PMNs. Bars represent the mean ± SEM. *N* = 5-6 independent experiments. Significance determined by linear mixed effects model: Survival ~ Treatment + (1 | Subject) + $H_2O_2$ Concentration. Intercept (92.893* ± 7.055), PMN Treatment (12.610* ± 3.284), $H_2O_2$ Concentration (-53.225* ± 4.940), *p<0.001.

**<u>Supplemental Table 1</u>: MIC of strains 130, 3X130, and H041.** Minimum inhibitory concentration (MIC) for the indicated antibiotics for 130, 3X130, and H041 was determined by E-test. 3X130 showed increased resistance to multiple antibiotics. H041 showed higher resistance to penicillin/cephalosporins, doxycycline, and vancomycin than 3X130, suggesting additional genetically encoded adaptations in the H041 lineage.

| | **MIC (µg/mL)** | | |
|---|---|---|---|
| **Antibiotic** | **130** | **3x130** | **H041** |
| Benz-Penicillin | 0.047-0.064 | 0.38 | 1.5-2.0 |
| Erythromycin | 0.094-0.125 | 1 | 1 |
| Cefixime | <0.10 | 1.0-1.5 | 4 |
| Ceftriaxone | <0.10 | 0.38-0.5 | 1.5 |
| Doxycycline | 0.19-0.38 | 1.0-1.5 | 1.5-2.0 |
| Vancomycin | 4.0-12.0 | 12.0-16.0 | 24 |

**<u>Supplemental Table 2:</u> Table of KEGG pathways enriched for Gc genes DE in Gc+PMN_1h vs Gc+PMN_0h comparisons.**

| KEGG Pathway ID | Pathway Name | Gene Found | Gene Pathway | Percentage | p value | p value adj |
|---|---|---|---|---|---|---|
| 220 | Arginine biosynthesis | 4 | 12 | 0.330 | 0.000 | 0.007 |
| 760 | Nicotinate and nicotinamide metabolism | 3 | 12 | 0.250 | 0.005 | 0.045 |
| 910 | Nitrogen metabolism | 5 | 6 | 0.830 | 0.000 | 0.000 |
| 920 | Sulfur metabolism | 3 | 7 | 0.430 | 0.000 | 0.007 |
| 1100 | Metabolic pathways | 26 | 369 | 0.070 | 0.000 | 0.004 |
| 1120 | Microbial metabolism in diverse environments | 11 | 86 | 0.130 | 0.002 | 0.023 |
| 1210 | 2-Oxocarboxylic acid metabolism | 6 | 20 | 0.300 | 0.000 | 0.004 |
| 2010 | ABC transporters | 7 | 31 | 0.230 | 0.000 | 0.007 |

**LEGENDS FOR DATASETS**

**Dataset S1:Complete overview of sequenced and mapped reads, and PanOCT table for Gc gene mapping**

Tab 1: Overview and statistics for human samples
Tab 2: Overview and statistics for 130 and 3X130 samples
Tab 3: Overview and statistics for H041 samples
Tab 4: RIN Scores
Tab 5: PanOCT table for converting 130, 3X130 and H041 gene locus tags into FA1090 NGO_xxxx locus tags

**Dataset S2: Raw and Normalized read counts for genes in each Gc variant**

Tab 1: Raw counts 130
Tab 2: VST counts 130
Tab 3: Raw Counts 3X130
Tab 4: VST counts 3X130
Tab 5: Raw Counts H041
Tab 6: VST Counts H041
Tab 7: Raw Counts Host/PMN
Tab 8: VST Counts Host/PMN

**Dataset S3: L2FC and significance for each individual comparison for Gc**

Tab 1: 130+PMN_0h vs 130+PMN_1h
Tab 2: 130+PMN_1h vs 130_1h
Tab 3: 3X130+PMN_0h vs 3X130+PMN_1h
Tab 4: 3X130+PMN_1h vs 3X130_1h
Tab 5: H041+PMN_0h vs H041+PMN_1h
Tab 6: H041+PMN_1h vs H041_1h
Tab 7: Contextualized Gc+PMN_1h vs Gc+PMN_0h
Tab 8: DE genes shared by 130, 3X130, and H041 for Gc+PMN_1h vs Gc+PMN_0h

**Dataset S4: L2FC and significance for each individual comparison for PMNs**

Tab 1: 130+PMN_0h vs 130+PMN_1h
Tab 2: 3X130+PMN_0h vs 3X130+PMN_1h
Tab 3: H041+PMN_0h vs H041+PMN_1h
Tab 4: PMN_0h vs PMN_1h
Tab 5: Time-Adjusted_PMN - 130+PMN_0h vs 130+PMN_1h
Tab 6: Time-Adjusted_PMN - 3X130+PMN_0h vs 3X130+PMN_1h
Tab 7: Time-Adjusted_PMN - H041+PMN_0h vs H041+PMN_1h
Tab 8: Time-Adjusted_PMN DE genes with average L2FC Gc+PMN_0h vs Gc+PMN_1h and average VST count for Gc+PMN_1h conditions.

**Dataset S5: IPA analysis results**

Tab 1: List of enriched Canonical Pathways and associated Time-Adjusted DE genes
Tab 2: List of enriched Molecular Functions and associated Time-Adjusted DE genes
Tab 3: List of enriched Molecular Functions subcategories and associated Time-Adjusted DE genes for each subcategory.

**Dataset S6: GSEA analysis results**

Tab 1: All GSEA datasets comparable to our input Time-Adjusted DE genes. Our top 20 expressed genes are highlighted in blue. Datasets that are referenced in the supplemental results/discussion are hyperlinked to their location in GSEA.
